# A lineage-specific Exo70 is required for receptor kinase-mediated immunity in barley

**DOI:** 10.1101/2021.12.19.473371

**Authors:** Samuel Holden, Molly Bergum, Phon Green, Jan Bettgenhaeuser, Inmaculada Hernández-Pinzón, Anupriya Thind, Shaun Clare, James M. Russell, Amelia Hubbard, Jodi Taylor, Matthew Smoker, Matthew Gardiner, Laura Civolani, Francesco Cosenza, Serena Rosignoli, Roxana Strugala, István Molnár, Hana Šimková, Jaroslav Doležel, Ulrich Schaffrath, Matthew Barrett, Silvio Salvi, Matthew J. Moscou

## Abstract

In the evolution of land plants, the plant immune system has experienced expansion in immune receptor and signaling pathways. Lineage-specific expansions have been observed in diverse gene families that are potentially involved in immunity, but lack causal association. Here, we show that *Rps8*-mediated resistance in barley to the fungal pathogen *Puccinia striiformis* f. sp. *tritici* (wheat stripe rust) is conferred by a genetic module: *LRR-RK* and *Exo70FX12*, which are together necessary and sufficient. The *Rps8* LRR-RK is the ortholog of rice extracellular immune receptor Xa21 and Exo70FX12 is a member of the Poales-specific Exo70FX clade. The Exo70FX clade emerged after the divergence of the Bromeliaceae and Poaceae, and comprises from 2 to 75 members in sequenced grasses. These results demonstrate the requirement of a lineage-specific Exo70FX12 in *Rps8* LRR-RK immunity and suggest that the Exo70FX clade may have evolved a specialized role in receptor kinase signaling.

## Introduction

Eukaryotes descend from a common ancestor that had diverse multimeric complexes that controlled transcription, translation, protein degradation, and secretion (Eme et al., 2017). As lineages diverged, these complexes experienced unique developmental, environmental, and non-self interactions that would shape their evolutionary trajectories. As sessile organisms, plant genomes have experienced a drastic expansion in signaling pathways that relay information to regulate biological processes. These include plant-specific protein kinases and membrane-bound receptor kinases (Lespinet et al., 2002), a proportion of which form essential components of the plant immune system. Throughout their evolution, plants have been exposed to diverse microbes, among which a subset are pathogenic. Recognition of pathogens by plants is mediated by two major classes of immune receptors, which are classified by the spatiotemporal properties of the interaction. The plant immune response requires pathogen detection, either at the cell boundary by membrane-localized extracellular receptors, or during infection within the cell by intracellular receptors (Dodds and Rathjen, 2010). This distinction is based on observed differences in both the signaling mechanisms and downstream responses between these interactions, however there is also clear interaction between these signaling pathways (Navarro et al., 2004; Tsuda and Katagiri, 2010; Thomma et al., 2011; Ngou et al., 2020; Yuan et al., 2021). Pathogens perturb plants through the synthesis and deployment of molecules such as toxins, catabolic enzymes and effectors which facilitate infection. Effectors are molecules generated by pathogens which manipulate their host to the benefit of the pathogen. This can include overcoming and suppressing immunity, directing the flow of nutrients to the pathogen, and altering the host to facilitate the pathogen’s life-cycle (Stergiopoulos and de Wit, 2009; Mukhtar et al., 2011; Lo Presti et al., 2015). Effectors are translocated to the plant cell and depending on the effector, can localize to the apoplast, the cytosol, or other subcellular compartments.

The majority of intracellular immune receptors belong to the nucleotide binding, leucine-rich repeat (NLR) class of receptor. The intracellular defense response is typically more rapid and intense than extracellular-based responses (Cui et al., 2015; Jones et al., 2016) and is associated with lineage-specific effector recognition. Extracellular recognition occurs at the boundary to the cell, generally via membrane-bound receptor proteins (RP) or receptor kinase proteins (RK), which are classified based on whether the receptor has an integrated kinase domain for downstream signaling (Zipfel and Oldroyd, 2017). These receptors are composed of an N-terminal exogenous binding domain, a transmembrane domain, and C-terminal cytoplasmic kinase domain in RK proteins. Extracellular immune receptors recognize non-self molecular patterns associated with microbial activity via direct binding to epitopes (Newman et al., 2013) such as the bacterial flagellin peptide fragment flg22 (Felix et al., 1999), the chitin oligomer N-acetylchitoheptose (Roby et al., 1987; Yamada et al., 1993) or plant-derived peptidoglycan cell wall fragments (Gust et al., 2007). In these examples, the recognized epitopes are conserved across a wide range of pathogens (e.g. fungal chitin), thus permitting a single receptor to provide immunity against a related group of pathogens. A subset of extracellular immune receptors recognize lineage specific epitopes, such as Xa21 (Ronald et al., 1992; Song et al., 1995; Pruitt et al., 2015) and Stb6 (Chartrain et al., 2005; Saintenac et al., 2018) that recognize peptides specific to *Xanthomonas* spp. and *Zymoseptoria tritici*, respectively.

Both classes of membrane-bound receptor proteins are further categorized by the exogenous receptor domain at the N-terminus of the protein. There are 14 common exogenous domains found in green plants, as well as various noncanonical integrations (Shiu and Bleecker, 2003; Shiu et al., 2004). Classes of ectodomains are also specialized towards particular classes of ligand. For example leucine-rich repeat (LRR) domains primarily associate with peptide-derived ligands (Kobe and Kajava, 2001), LysM domains with carbohydrate ligands (Buist et al., 2008), and lectins with lipopolysaccharides (Lannoo and Van Damme, 2014). LRR-RK proteins are the largest single class of RK proteins in plants, and LRR domains have been identified in all five kingdoms of life (Morillo and Tax, 2006; Fischer et al., 2016). The LRR domain is generally composed of up to 30 leucine-rich repeats which vary from 20 to 29 amino acids, and a small cap at the N-terminus. A strong phylogenetic signature facilitates the classification of RK proteins based on the protein kinase domain. The RK/Pelle encoding genes of *Arabidopsis thaliana* and rice have been classified into over 60 families, of which 15 families were LRR-RK-specific (Shiu and Bleecker, 2003; Shiu et al., 2004; Lehti-Shiu et al., 2009). The LRR-XII subfamily is dramatically expanded in rice (>100 members) compared to *A. thaliana* (10 members) and contains several of the most studied plant defense LRR-RK genes, including *FLS2* (Gomez-Gomez and Boller, 2000) and *EFR* (Zipfel et al., 2006) in dicotyledonous plants and *Xa21* in the Oryzoideae (Song et al., 1995). FLS2 recognizes a 22 amino acid epitope derived from bacterial flg22 (Chinchilla et al., 2006) and EFR recognizes an 18 amino acid peptide fragment (elf18) of the bacterial elongation factor Ef-Tu, whereas Xa21 recognizes a 20 amino acid tyrosinated region (RaxX21-sY) derived from RaxX, a type-1 secreted peptide of 61 residues that is hypothesized to have effector activity as a mimic of plant growth hormones (Pruitt et al., 2015; Pruitt et al., 2017). Extensive work in *A. thaliana* has shown that after ligand perception and binding, the protein-ligand complex is able to form a heterodimer with the LRR-II type RK BAK1 (Chinchilla et al., 2007; Schulze et al., 2010) which interacts with both the extracellular bound-ligand and the intracellular membrane-bound components of FLS2 (Sun et al., 2013). EFR signals via the co-receptor BAK1, with similar downstream components as FLS2 (Zipfel et al., 2006; Meng and Zhang, 2013), and AtBAK1 paralogue OsSERK2 is required for Xa21 signaling, suggesting an ancestral requirement of the LRR-XII subfamily for this co-receptor (Chen et al., 2014).

The *Pucciniales* are an order of obligate biotroph fungal pathogens, and causal agents of rust diseases on a wide variety of host plants (McTaggart et al., 2016; Chen and Kang, 2017). Particularly relevant to the cereal crops are *Puccinia striiformis* (stripe rust), *Puccinia triticina* (leaf rust), and *Puccinia graminis* (stem rust) (Roelfs and Bushnell, 1985). *P. striiformis* is found in all major wheat growing areas and impacts global production by over 2% (Savary et al., 2019). *P. striiformis* has a complex five stage lifecycle, although only the uredinial stage directly impacts cereal production. *P. striiformis* exhibits host specialization, *formae speciales*, such that isolates derived from wheat (*P. striiformis* f. sp. *tritici*; *Pst*) do not routinely infect barley, whereas isolates derived from barley (*P. striiformis* f. sp. *hordei*) do not routinely infect wheat (Eriksson, 1894, 1898). This host specialization shows evidence of durability, as *Pst* has been present for over 60 years in Australia, but has not undergone a host jump on to barley, despite being planted in similar regions with wheat (Wellings, 2007). Screening of diverse barley accessions has found that *Pst* is capable of infecting and reproducing on a small subset of barley accessions, primarily landraces and wild barley, as well as the hypersusceptible accession SusPtrit (Yeo et al., 2014; Dawson et al., 2015; Niks et al., 2015; Li et al., 2016; Bettgenhaeuser et al., 2021). Using a diverse panel of barley accessions, we previously identified three *R* gene loci designated *Rps6, Rps7*, and *Rps8* as contributing to the non-adapted status of barley to *Pst* (Bettgenhaeuser et al., 2021). *Rps8* was mapped to the long-arm of chromosome 4H using a mapping population derived from SusPtrit x Golden Promise (Yeo et al., 2014; Bettgenhaeuser et al., 2021). In this work, we fine mapped *Rps8* to a 936 kb locus on chromosome 4H, which encompasses as a presence/absence variation across barley accessions. Forward genetic screens and transgenic complementation demonstrate that resistance is conferred by a genetic module including an *LRR-RK* and *Exo70*, which are necessary and together, sufficient for *Rps8*-mediated resistance.

## Results

### Fine mapping of *Rps8* resolved the locus to a 936 kb region on chromosome 4H

*Rps8* is located on the long arm of chromosome 4H and prevents the development of pustules of wheat stripe rust, but not colonization (**Figure 1a**). Inheritance of *Rps8* in an F_2_ population generated by crossing SxGP DH-21 (*rps8*) and SxGP DH-103 (*Rps8*) from the doubled-haploid SusPtrit x Golden Promise population (Yeo et al., 2014) found clear segregation for a single dominant gene (**Figure 1bc**). A high-resolution recombination screen was performed using markers generated in the *Rps8* region, identifying 127 recombinants among 9,216 gametes that resolved *Rps8* to a 0.5 cM genetic interval. This interval corresponds to a 936 kb physical interval in the genome of the reference accession Morex, which contains the *Rps8* haplotype (Mascher et al., 2021) (**Figure 1d**). Markers co-segregating with *Rps8* localize within a 664 kb physical interval. Nine recombination events were identified proximal to the markers co-segregating with *Rps8* and fifteen in the distal region (**Figure 2a**). Analysis of genes in the *Rps8* region in Morex found 17 annotated protein encoding genes: a DUF4371 protein, a DUF1997 protein, an Armadillo-repeat protein (ARM), a Exocyst subunit Exo70 (Exo70), a transmembrane receptor protein kinase (LRR-RK), two BTB/POZ protein, a putatively fragmented NLR (CC-NB and LRR-DDE), a zinc-finger transcription factor (ZnF), a Myb transcription factor (Myb), several unknown proteins, and several annotated transposon-derived proteins (**Figure 2a; Supplemental Table 1)**. To evaluate the candidacy of individual genes in the interval, genetic, genomic, and transcriptomic data were evaluated for every gene. To identify genes expressed in leaf tissue, we used a previously generated RNAseq tissue atlas derived from Morex (IBGSC, 2012). Twelve genes were expressed in at least one tissue type and eight genes were found to be expressed in leaf at detectable levels (**Supplemental Table 1**). The fragmented NLR was found to be expressed in roots but not leaves (**Supplemental Figure 1**). Aligned RNAseq data confirmed that the NLR did not have an intact in-frame open reading frame. In leaf, non-repetitive expressed genes include the *DUF1997, ARM, Exo70, LRR-RK, ZnF* and *Myb* transcription factors (**Supplemental Figure 1**). To understand the expression of these genes in haplotypes with and without *Rps8*, we leveraged previously generated genetic data that established the presence or absence of *Rps8* in diverse barley accessions and leaf RNAseq from these accessions (**Figure 3a**) (Dawson et al., 2016; Maekawa et al., 2019; Bettgenhaeuser et al., 2021). While the genes *PPR* (670.1), *ARM* (isoforms 700.1 and 700.2), and a transposon Gag-Pol polyprotein (770.1) were expressed, no specific pattern was observed in *Rps8* and *rps8* haplotypes (**Figure 3a**). In contrast, all accessions carrying *Rps8* were found to express *Exo70* and *LRR-RK*, whereas the majority of accessions lacking *Rps8* did not express these genes (**Figure 3bc**). Exceptions included the barley accession Heils Franken and WBDC247. These results indicate that for the majority of accessions, *Exo70* and *LRR-RK* have an expression level polymorphism that is correlated with the presence of a functional *Rps8* haplotype.

**Figure 1.**
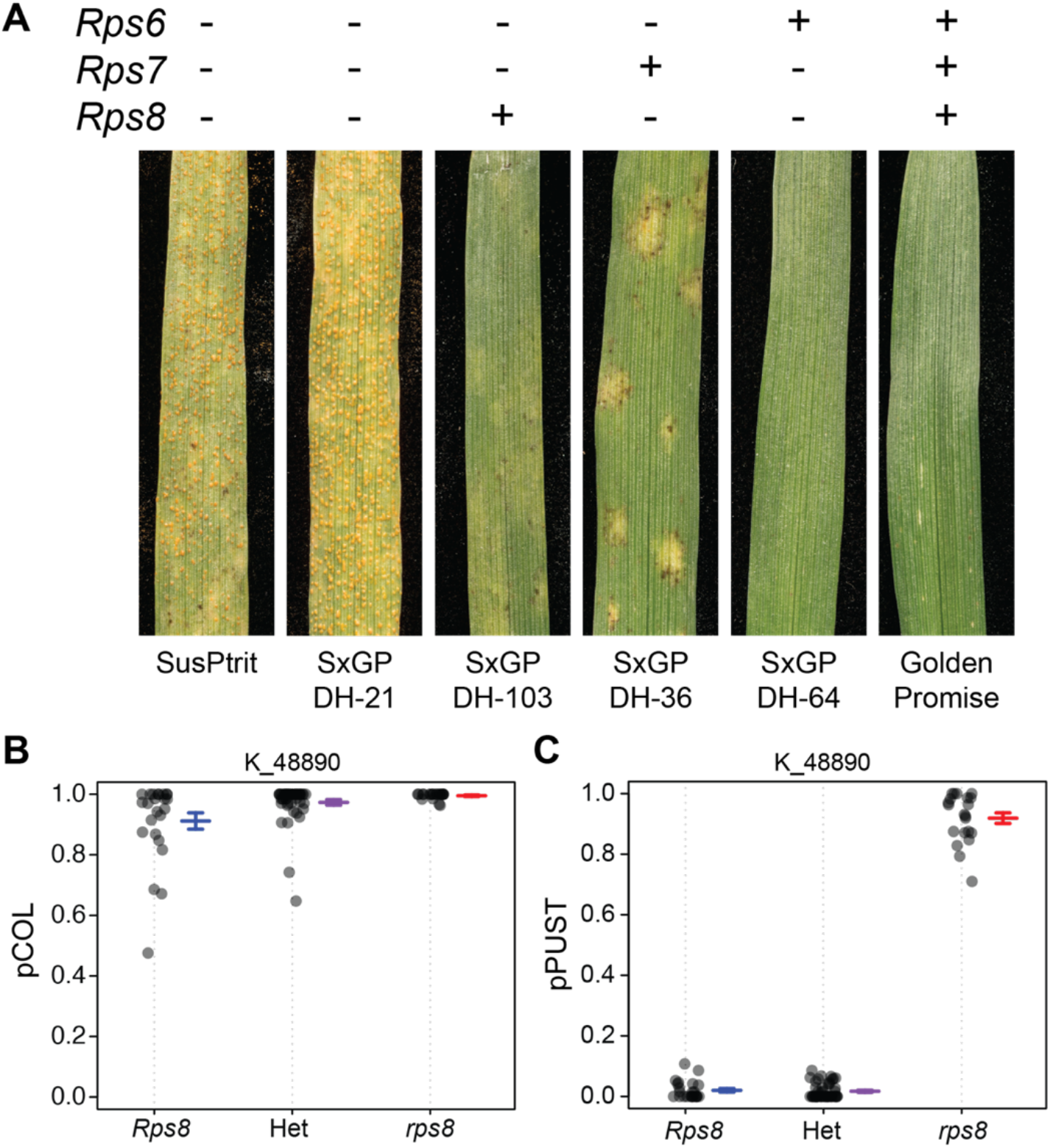
*Rps8* prevents pustule formation, but not colonization of wheat stripe rust (*Puccinia striiformis* f. sp. *tritici*) in barley. (A) Presence (+) or absence (-) of the resistance genes *Rps6, Rps7*, and *Rps8* in representative accessions in the Golden Promise x SusPtrit doubled-haploid population and phenotype from first leaf inoculated with *Pst* isolate 16/035 at 14 dpi (Bettgenhaeuser et al., 2021). (B) and (C) Colonization (pCOL) and pustule formation (pPUST) were assessed in a SxGP DH-21 (*rps6 rps7 rps8*) x SxGP DH-103 (*rps6 rps7 Rps8*) F_2_ population inoculated with *Pst* isolate 16/035 (N=94). Allelic states include homozygous *Rps8* (*Rps8*), heterozygous (Het), and homozygous *rps8* (*rps8*). *Rps8* co-segregates with the KASP marker K_48890.

**Figure 2.**
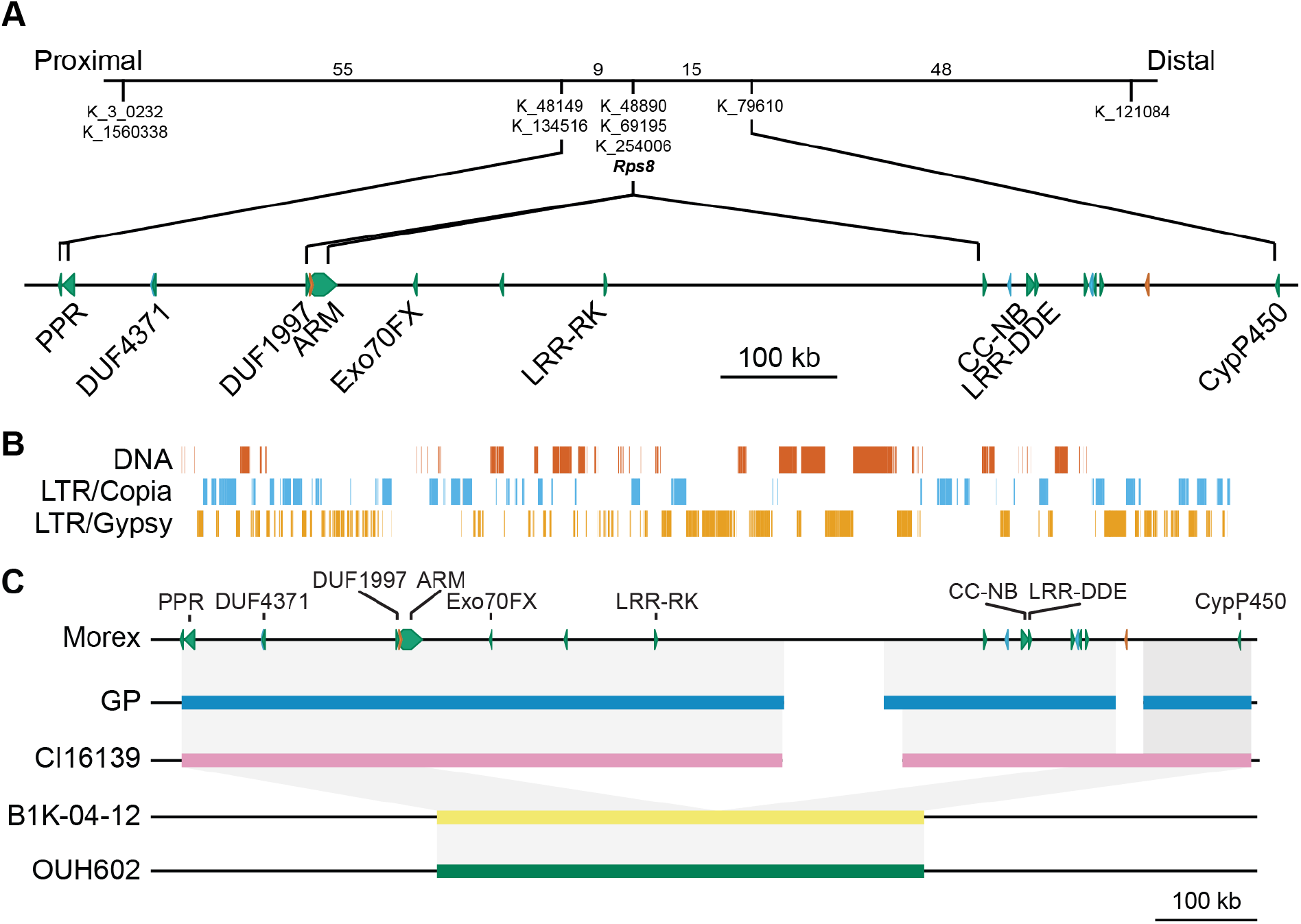
*Rps8* maps to a 900 kb region on chromosome 4H and is associated with the presence of a 546 kb insertion. (A) High-resolution recombination screen was resolved *Rps8* to a 0.5 cM genetic interval based on 9,216 gametes. The physical interval corresponds to 936 kb in the genome of the reference accession Morex (*Rps8*) (Mascher et al., 2021). The seventeen annotated protein encoding genes include a DUF4371 protein, a DUF1997 protein, a Armadillo-repeat protein (ARM), a Exocyst subunit Exo70 (Exo70), a transmembrane receptor protein kinase (LRR-RK), two BTB/POZ protein, a fragmented NLR (CC-NB and LRR-DDE), a zinc-finger transcription factor (ZnF), a Myb transcription factor (Myb), several unknown proteins, and several annotated transposon-derived proteins. Numbers in the genetic interval are the number of identified recombinants. (B) Repetitive content of the *Rps8* region. A tandem array of the 5’ repeat region of a CACTA DNA transposon interrupted by two head-to-head long-terminal repeat (LTR) retrotransposons is located in the center of the region. (C) Barley accessions carrying *Rps8* are highly conserved. Genomes of Morex (*Rps8*) (Mascher et al., 2021), Golden Promise (*Rps8*) (Schreiber et al., 2020), the wild barley accessions B1K-04-12 (Jayakodi et al., 2020) and OUH602 (*rps8*) (Sato et al., 2021), and a *de novo* sequenced chromosome 4H of accession CI16139 (*Rps8*). A large (190 kb), repetitive region identified in Morex was a breakpoint in the assembly of the *Rps8* locus in CI16139 and Golden Promise. The wild barley accession B1K-04-12 is collapsed relative to the other genomes, indicating that a 546 kb InDel encompasses six genes: *Exo70, LRR-RK, CC-NB, LRR-DDE, CC-TM*, and transposon.

**Figure 3.**
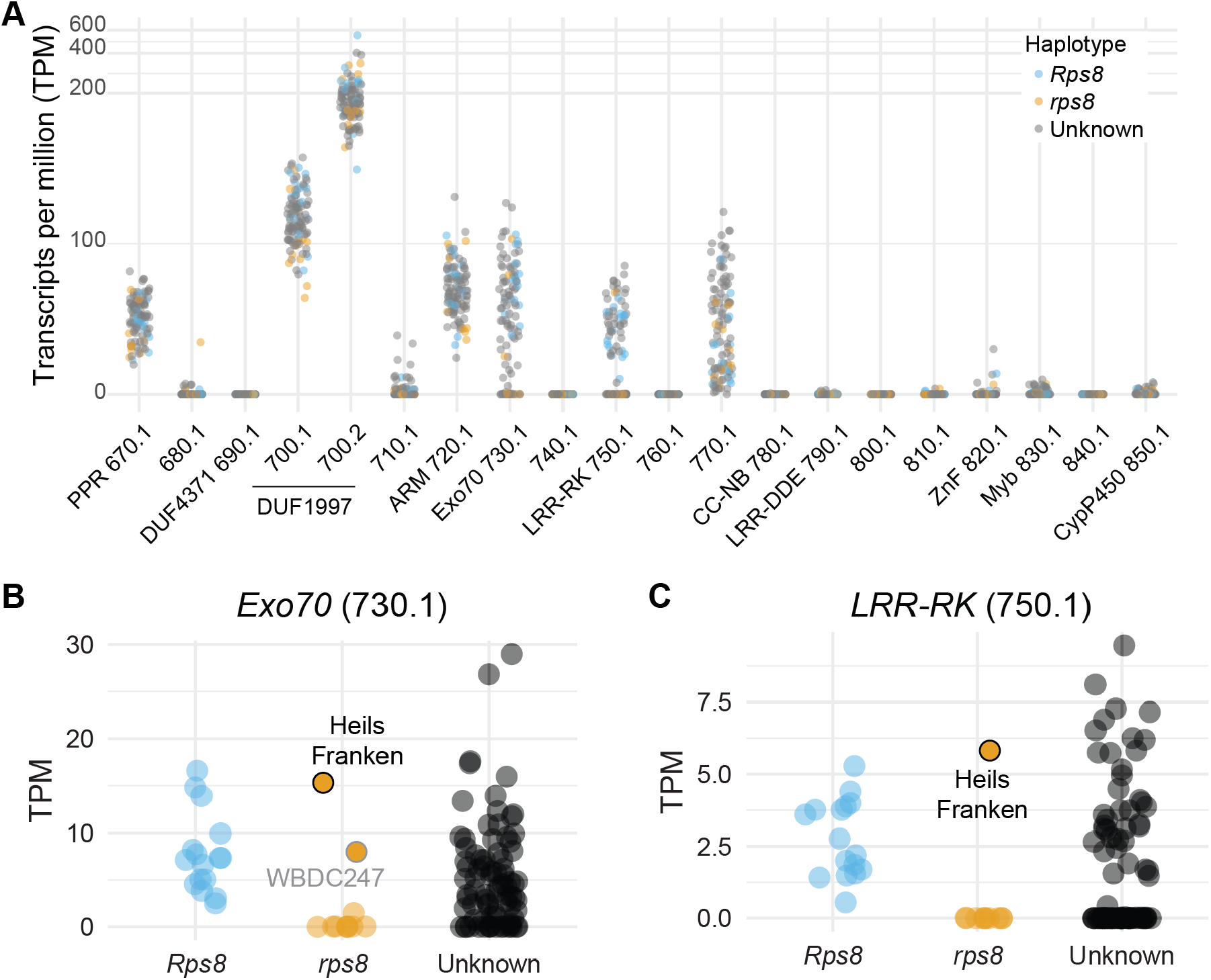
Expression level polymorphism in *Rps8 LRR-RK* and *Exo70* is correlated with the presence of a functional *Rps8* haplotype. (A) Leaf expression level of genes in 109 barley accessions in the *Rps8* region. Identifiers (x-axis) correspond to the suffix of genes within the *Rps8* locus based on the longer identifier HORVU.MOREX.r3.4H0407xxx. The y-axis shows expression level based on transcripts per million (TPM). (B) and (C) Expression level polymorphism in *Rps8 LRR-RK* and *Exo70* is correlated with the presence of a functional *Rps8* haplotype. Association of presence or absence of *Rps8* in diverse barley accessions and leaf RNAseq for *Rps8 LRR-RK* (*HORVU*.*MOREX*.*r3*.*4H0407750*.*1*) and *Exo70* (*HORVU*.*MOREX*.*r3*.*4H0407730*.*1*). The expression level polymorphism in *Rps8 Exo70* and *LRR-RK* is predictive for *Rps8*-mediated resistance except in the barley accessions Heils Franken and WBDC247.

### *Rps8* encompasses a 546 kb InDel

We hypothesized that the expression polymorphism in *Exo70* and *LRR-RK* was due to structural variation in the *Rps8* locus between resistant and susceptible haplotypes. To test this hypothesis, we compared the genomic regions encompassing *Rps8* in Morex (*Rps8*) (Mascher et al., 2021), Golden Promise (*Rps8*) (Schreiber et al., 2020), the wild barley accessions B1K-04-12 (Jayakodi et al., 2020) and OUH602 (*rps8*) (Sato et al., 2021), and a *de novo* sequenced chromosome 4H of accession CI 16139 (*Rps8*) using Chicago-based library and chromosome flow sorting (Thind et al., 2017). The sequence identity between the three *Rps8* haplotypes (Morex, Golden Promise, CI 16139) was extremely high, however a large (190 kb), repetitive region identified in Morex was a breakpoint in the assembly of the *Rps8* locus in CI 16139 and Golden Promise (**Figure 2bc**). This region is defined by a tandem array of the 5’ repeat region of a CACTA DNA transposon interrupted by two head-to-head long-terminal repeat (LTR) retrotransposons (**Figure 2b**). Comparing these three haplotypes to the wild barley accessions B1K-04-12 and OUH602 found a 546 kb InDel that encompasses six genes: *Exo70, LRR-RK, CC-NB, LRR-DDE, CC-TM*, and transposon. (**Figure 2c)**. A region including the gene *HORVU*.*MOREX*.*r3*.*4HG0407760* (BTB/POZ domain protein) is present in the B1K-04-12 and OUH602 haplotypes at the junction of the InDel. These results show that the underlying variation observed in accessions lacking *Rps8* involve a loss of the interval.

### Natural variation identifies a loss-of-function mutant in *Exo70*

The presence of a 546 kb InDel limited our ability to resolve the fine genetic structure of the *Rps8* region. Therefore, an alternative strategy to map-based cloning is required to dissect the *Rps8* locus. Our next approach was to interrogate other sources of variation to evaluate candidate genes underlying *Rps8*. RNAseq data from a panel of 109 barley accessions were aligned to the *Rps8* region (**Supplemental Table 2**). Expression of *LRR-RK* and *Exo70* was observed in 56 accessions and absent in 53 accessions. Analysis of protein sequence found 15 LRR-RK and 10 Exo70 variants, which together represent 19 unique haplotypes expressing *LRR-RK* and *Exo70*. The majority of accessions (N=33) expressing *LRR-RK* and *Exo70* had identical protein sequence with the Morex haplotype. All other haplotypes were identified in wild barley except for the landrace Heils Franken, which carries a single non-synonymous polymorphism in *Exo70* that causes an E271K substitution.

While the barley accession Heils Franken is resistant to *Pst*, the underlying genetic architecture for this resistance is unknown. To determine whether resistance is conferred by *Rps8*, Heils Franken x Manchuria F_2_ and BC_1_ populations were generated and inoculated with *Pst* isolate 08/21. In the Heils Franken x Manchuria F_2_ population (N=94), the majority of the phenotypic variation was explained by a single locus on chromosome 1H (PVE=95.2%) for macroscopic chlorosis and microscopic colonization phenotypes. This locus coincides with *Mla* (=*Rps7*), suggesting that Heils Franken may carry a functional *Mla*/*Rps7* allele to *Pst*. There was insufficient power to detect the presence of *Rps8* in this population, as only seven individuals exhibited pustule formation greater than 0.5 (**Supplemental Figure 2**). In contrast, the Heils Franken x Manchuria BC_1_ population (N=94) and derived progeny exhibited considerable variation for colonization and pustule formation to *Pst* (**Supplemental Figure 3**). Interval mapping identified a single major-effect locus on chromosome 1H providing resistance, a major effect QTL on chromosome 5H, and a minor effect QTL on chromosome 4H that is distinct from *Rps8* (**Figure 4a**). The chromosome 1H QTL, previously identified in the Heils Franken x Manchuria F_2_ population, controls a significant proportion of the phenotypic variation for chlorosis (PVE=34.3%), colonization (PVE=41.6%), infection (PVE=18.7%), and pustule formation (PVE=17.5%). In contrast, no significant linkage is observed for *Rps8*, indicating that the Heils Franken haplotype does not provide resistance to *Pst* (**Figure 4b**).

**Figure 4.**
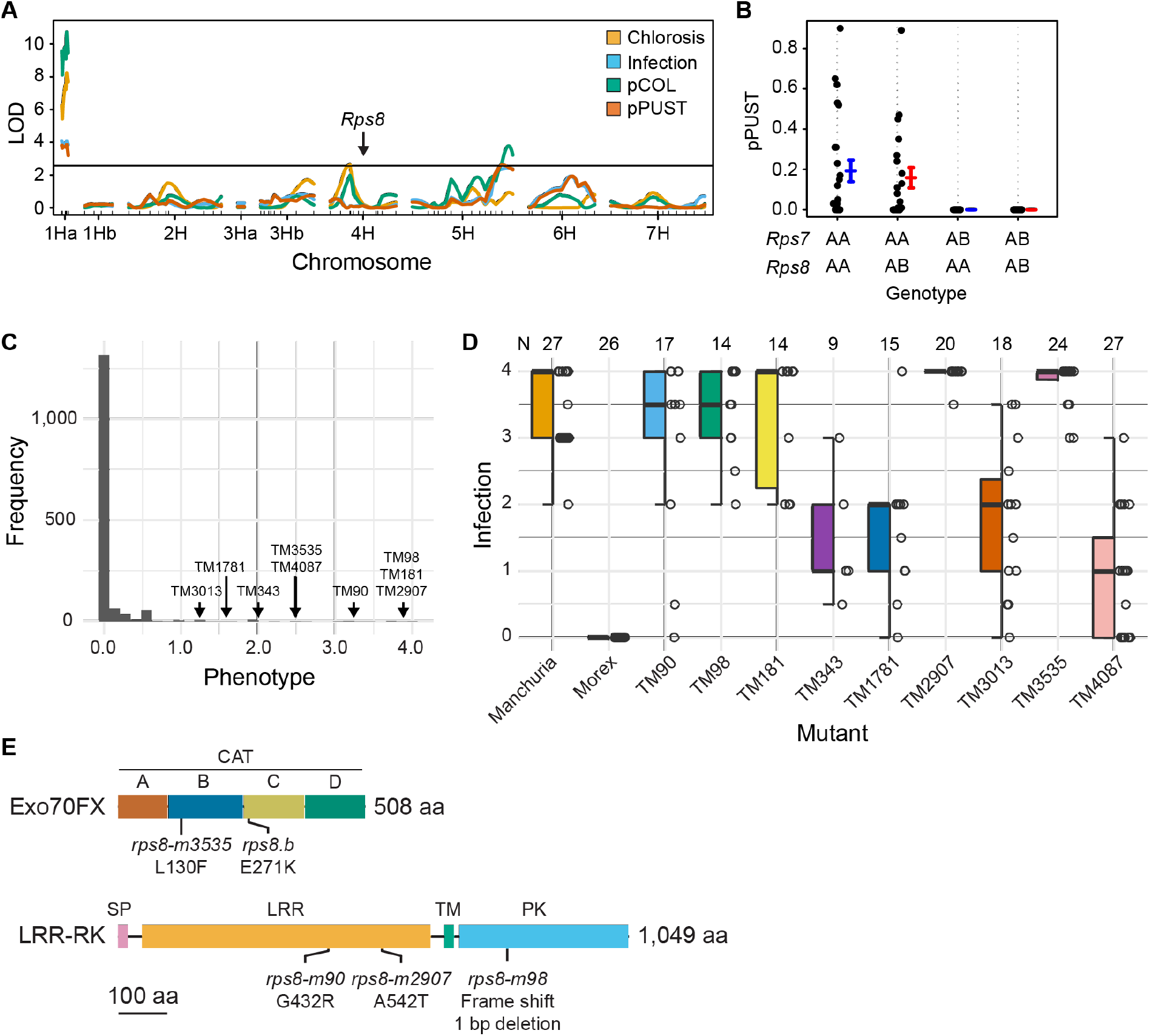
Natural and induced variation identifies loss-of-function mutations in the *Exo70* and *LRR-RK* at *Rps8*. (A) Quantitative trait locus analysis of *Pst* isolate 16/035 resistance in the Manchuria x Heils Franken BC_1_ population. QTLs conferring resistance were identified on 1H, 4H, and 5H, but none overlapped with the *Rps8* locus. (B) Phenotype x genotype plot of the *Rps7* and *Rps8* loci based on the Manchuria x Heils Franken BC_1_ population from (A). Strong linkage is observed for *Rps7*, but not *Rps8* for pPUST (pustule formation). (C) Histogram of infection phenotypes (macroscopic pustule formation) from a forward genetic screen using sodium azide mutagenized Morex (TM population) M_6_ families inoculated with *Pst* isolate 16/035. Among 1,526 M_6_ families, a total of 35 putative mutants were identified with infection phenotypes of 1.0 or higher based on an average of eight seedlings in a single experiment. (D) Infection phenotypes of confirmed mutants using *Pst* isolate 16/035. Panel shows boxplots, individual data points, and total number of evaluated plants (N) over three biological replicates. (E) Position of missense and nonsense mutations from sodium azide induced and natural variation in barley accession Heils Franken in *Rps8* Exo70 and LRR-RK. Sub-domain structure of Exo70 is based on predictions from yeast. Domain structure of LRR-RK is based on InterProScan predictions.

To associate the mutation in *Exo70* with loss of *Rps8*-mediated resistance, it is essential to show that no other variation exists within the *Rps8* locus in Heils Franken as compared to Morex. We performed whole genome sequencing of Heils Franken. Alignment and variant calling found that the only variant in annotated protein encoding genes in the *Rps8* region was the previously identified non-synonymous polymorphism in *Exo70* creating a E271K modification. This suggests that the *Exo70* may be required for *Rps8*-mediated resistance.

### A forward genetic screen identifies loss-of-function mutations in *LRR-RK* and *Exo70*

While natural variation provided evidence that *Exo70* is likely required for *Rps8*-mediated resistance, the remaining variation was insufficient to establish causal relationships. We initiated a forward genetic screen to identify genes underpinning *Rps8*-mediated resistance that made use of a mutant population that had previously been generated using the reference accession Morex (TM population) (Talamè et al., 2008), which harbors *Rps8* in isolation from other loci providing *Pst* resistance. The TM population was generated using sodium azide and has been advanced to the M_6_ stage. We screened 1,526 M_6_ families with *Pst* isolate 16/035 and identified 37 putative mutants with an infection phenotype (area showing pustule formation) of 1.0 or higher (**Figure 4c; Supplemental Table 3**). Single seed descent of putative mutants and subsequent re-evaluation with *Pst* isolate 16/035 confirmed that nine mutants were susceptible to *Pst* (**Figure 4d**).

To identify polymorphisms in genes in the *Rps8* locus, we performed RNAseq on leaf tissue from these nine mutants. Alignment of RNAseq found three independent mutations in the *LRR-RK* CDS with missense mutations in *rps8-TM90* (CDS: G1409A, protein G432R) and *rps8-TM2907* (CDS: G1624A, protein A542T) in the LRR encoding region, and *rps8-TM98* carrying a G deletion at 2,504 bp in the kinase encoding region that causes a frame shift and early stop codon (**Figure 4e**). The mutant line *rps8-TM3535* had a mutation in the *Exo70* CDS, exhibiting a C to T transition at 388 bp that leads to a missense mutation (L130F) (**Figure 4e**). Analysis of RNAseq data from five additional mutant lines did not identify any variation in genes at the *Rps8* locus, indicating that these mutations likely occur in other genes that are *Required for* Rps8*-mediated resistance* (*Rsr*). To confirm the genetic relationship of *rps8* mutants, pairwise crosses were performed between a subset of mutants at the *Rps8* locus and F_1_ progeny inoculated with *Pst* isolate 16/035. The mutants TM90 and TM98 were found to belong to the same complementation group, whereas TM3535 was in an independent complementation group (**Supplemental Figure 4**). These results confirm the requirement of both *LRR-RK* and *Exo70* for *Rps8*-mediated resistance.

### *LRR-RK* and *Exo70* are necessary and sufficient to confer *Rps8*-mediated resistance

We hypothesized that transgenic complementation using either *LRR-RK* or *Exo70* would be insufficient to confer *Rps8*-mediated resistance and that instead would require the presence of both genes. Our initial experiment was to determine whether stable transformation of *Exo70* is sufficient to confer *Rps8*-mediated resistance. We transformed the wheat stripe rust susceptible accession SxGP DH-47 using *Agrobacterium*-based transformation with a T-DNA construct that included the native genomic region encompassing *Exo70* (**Supplemental Figure 5**) (Hensel and Kumlehn, 2004). Eight independent single copy transgenic lines were generated and inoculated with *Pst* isolate 16/035. No specific association was observed in T_1_ transgenic families with the presence/absence of the T-DNA and segregating phenotypes (**Supplemental Figure 6**). This suggested that the *Exo70* is insufficient to confer *Rps8*-mediated resistance. To confirm that the *Exo70* transgene was functional, we crossed homozygous T_2_ individuals for the T_1_-10b, T_1_-7b, and T_1_-14a transformant lineages with the mutant TM3535 which carries a single non-synonymous mutation in *Exo70*. While homozygous T_2_ and T_3_ individuals from the single copy transgene lineages T_1_-10b, T_1_-7b, and T_1_-14a were susceptible, all F_1_ progeny derived from crosses with TM3535 were resistant to *Pst* isolate 16/035 (**Figure 5a**). Therefore, *Exo70* is sufficient to complement the mutant TM3535, but insufficient to confer *Rps8*-mediated resistance when transformed in SxGP DH-47.

**Figure 5.**
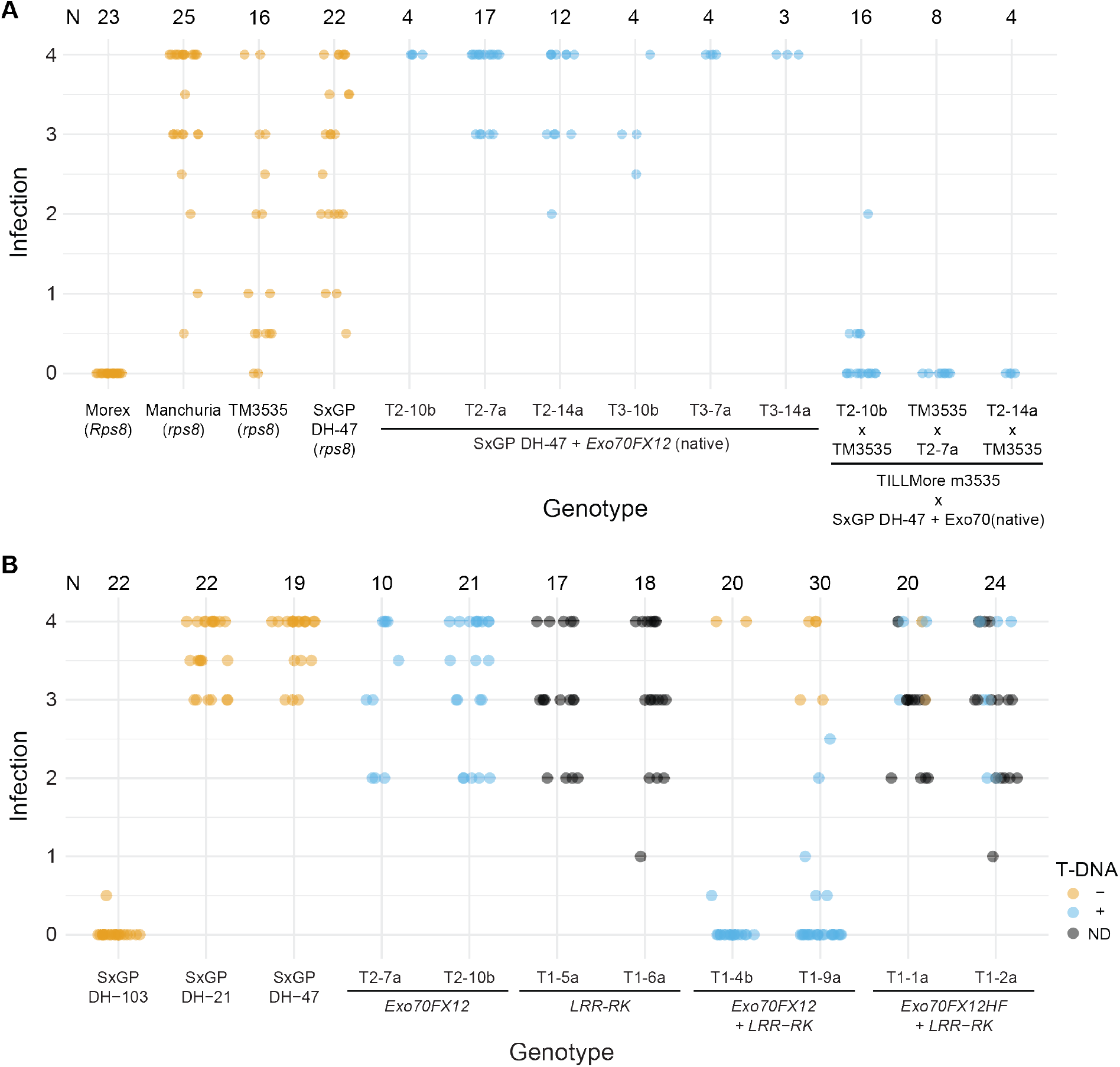
*LRR-RK* and *Exo70FX12* are sufficient to confer *Rps8*-mediated resistance. (A) Infection phenotypes for three independent single copy advanced T_2_ and T_3_ homozygous transformants of barley accession SxGP DH-47 expressing *Exo70FX12* under its native promoter and terminator, F_1_ based on a cross of transgenic *Exo70FX12* lines and the TM3535 *Exo70FX12* mutant, and controls Morex (*Rps8*), Manchuria (*rps8*), TM3535 (*rps8-TM3535*), and SxGP DH-47 (transformable, susceptible line (*rps8*)). (B) Infection phenotypes for independent single copy transformants of barley accession SxGP DH-47 expressing *Exo70FX12, LRR-RK, Exo70FX12* and *LRR-RK*, and *Exo70FX12HF* (Heils Franken allele) and *LRR-RK*, under their native promoters and terminators. Presence (blue) or absence (orange) of the T-DNA was determined using a qPCR-based assay. When not determined (ND), data points are in black. For both panels, inoculations were performed using *Pst* isolate 16/035 and scored at 14 days post inoculation, N shows the number of evaluated seedlings, and each panel represents a single experiment.

To test if expression of both *LRR-RK* and *Exo70* is required to confer *Rps8*-mediated resistance, we transformed SxGP DH-47 with T-DNA constructs that natively express *LRR-RK* individually, both *LRR-RK* and *Exo70*, and the *LRR-RK* with the *Exo70* allele from Heils Franken (**Supplemental Figure 5**). T_1_ families containing single T-DNA inserts, T_2_ families homozygous for natively expressed *Exo70*, and controls (Manchuria, Morex, SxGP DH-47) were inoculated with *Pst* isolate 16/035 (**Figure 5b**). T_1_ families carrying the *LRR-RK* expressed under its native promoter and T_2_ families homozygous for the *Exo70* expressed under its native promoter were susceptible to *Pst* isolate 16/035 (**Figure 5b**). In contrast, T_1_ families carrying a T-DNA containing both *LRR-RK* and *Exo70* under their native promoters conferred resistance that co-segregated with the presence of the T-DNA, whereas T_1_ families carrying *LRR-RK* and *Exo70* allele from Heils Franken were susceptible (**Figure 5b**). An extended set of transgenic T_1_ families predominantly supported these observations, although two transgenic families (T1-3b and T1-7a) expressing LRR-RK exhibited partial resistance, although these may have escaped inoculation or represent lines with higher levels of expression of the *LRR-RK* (**Supplemental Figure 7**). Collectively, these results demonstrate that together, the *LRR-RK* and *Exo70* are necessary and sufficient to confer complete *Rps8*-mediated resistance.

### The *LRR-RK* at *Rps8* belongs to the LRR-XII subfamily of RK and is the ortholog of *Xa21*

LRR-RK proteins are the predominant class of extracellular recognition receptor (Morillo and Tax, 2006; Fischer et al., 2016). *Rps8 LRR-RK* encodes a 1,080 amino acid protein with a domain structure of signal peptide (1-16 aa), 24 imperfect 24 amino acid LRRs (74-652 aa), transmembrane (670-695 aa), juxtamembrane (702-726 aa), and protein kinase (727-1,038 aa) (**Figure 4e, Supplemental Figure 8**). Using an HMM-based classification strategy developed by Lehti-Shiu and Shiu (2012), we found that the *Rps8* LRR-RK belongs to the LRR-XII subfamily. The LRR-XII subfamily includes the membrane immune receptors *Xa21* (Song et al., 1995), *FLS2* (Gomez-Gomez and Boller, 2000), and *EFR* (Zipfel et al., 2006). To ascertain the relationship of *Rps8* LRR-RK relative to other LRR-XII subfamily members, we constructed a phylogenetic tree based on full-length protein sequence from LRR-XII members from eight grass species (**Figure 6a; Supplemental Table 4**). *Rps8* LRR-RK was found in the same subclade as Xa21, which included RK from *B. distachyon* (9 RKs), *H. vulgare* (2 RKs), *O. sativa* (8 RKs), *Olyra latifolia* (3 RKs), *S. italica* (5 RKs), *S. bicolor* (1 RK), and *T. aestivum* (13 RKs) (**Supplemental Table 5**). Further phylogenetic analysis using this subclade, along with putative homologs in 24 additional grass species, found that *Rps8* LRR-RK and Xa21 are orthologs (**Figure 6b; Supplemental Tables 5 and 6**). Extensive phylogenetic and functional characterization has been performed on the putative orthologs of *Xa21* in the grasses (Tan et al., 2011; Cantu et al., 2013; Yang et al., 2013; Liu et al., 2016; Wang et al., 2019). In agreement with previous phylogenetic analysis, the wheat RK Xa21-like1 (TaXa21L1), Xa21-like2 (TaXa21L2), and Xa21-like3 (TaXa21L3) correspond to TraesCS4D01G352700.1, TraesCS2D01G585500.1, and TraesCS5A01G285000.1 (RefSeq2.0) (Cantu et al., 2013), and group distinct from the Xa21 subclade. In contrast, the RK encoding gene designated *TaXa21* is within the Xa21 subclade and is the wheat ortholog of *Rps8* LRR-RK (Wang et al., 2019). *TaXa21* was identified based on induced gene expression after treatment with high temperature and was associated with a quantitative reduction in *Pst* uredinia production in a temperature-dependent manner based on virus-based silencing. While no further genetic or functional characterization were performed, these results indicate that the orthologs of *Xa21* in barley and wheat both contribute to immunity to *Pst*.

**Figure 6.**
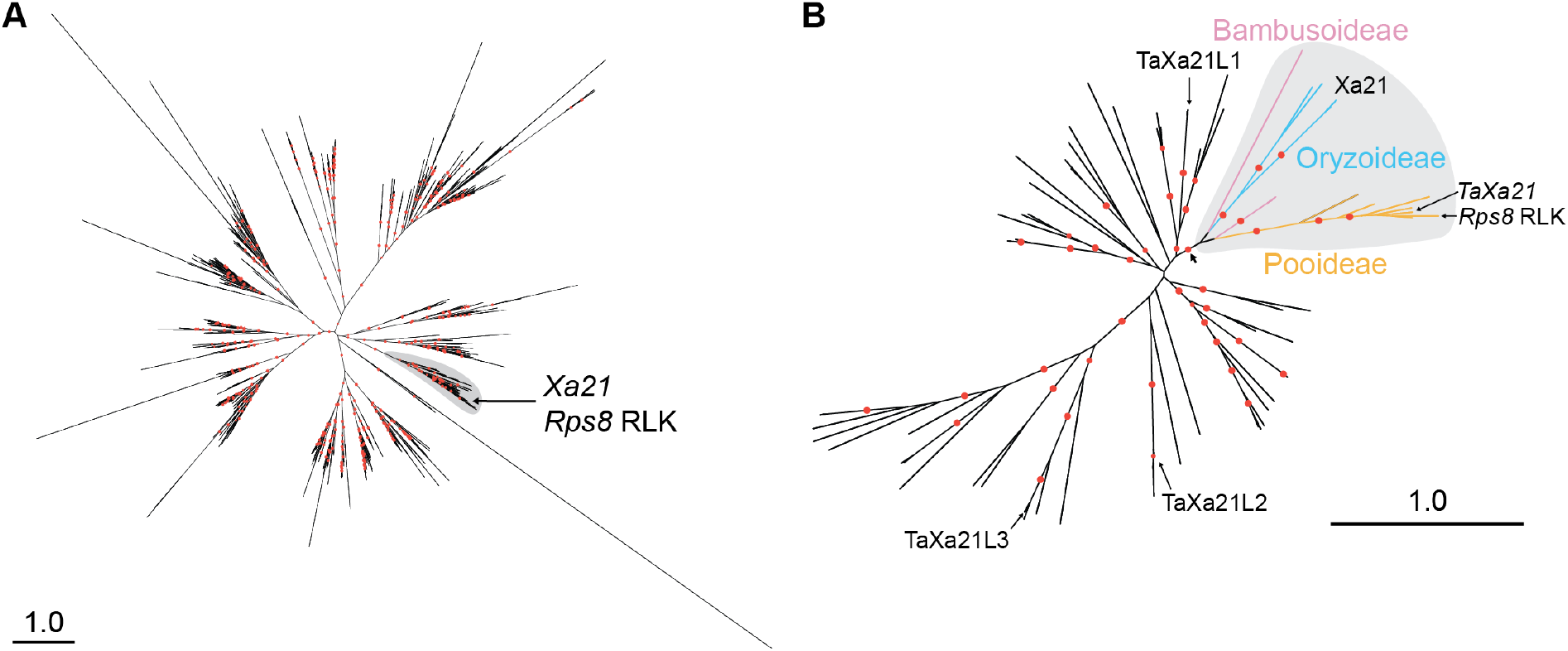
*Rps8 LRR-RK* belongs to the LRR-XII subfamily of RK and is the ortholog of *Xa21*. (A) Maximum likelihood unrooted phylogenetic tree based on 959 full length RK LRR-XII subfamily proteins from barley (*Hordeum vulgare*), wheat (*Triticum aestivum*), purple false brome (*Brachypodium distachyon*), rice (*Oryza sativa*), carrycillo (*Olyra latifolia*), maize (*Zea mays*), foxtail millet (*Setaria italica*), and sorghum (*Sorghum bicolor*). The Xa21/*Rps8* RK subclade is defined by the labelled grey region. Red dots indicate bootstrap support greater than 80%. (B) Maximum likelihood unrooted phylogenetic tree based on 69 full length RK proteins from the Xa21/*Rps8* RK clade and putative Xa21 homologs from 24 additional Poaceae species (**Supplemental Table 6**). Color coding on the subtree including Xa21 indicates the tribe origin of branches (Bambusoideae [pink], Oryzoideae [blue], and Pooideae [orange]). Red dots indicate bootstrap support greater than 80%. The basal branch on the Xa21 subtree indicated by an arrow is supported at 95%. Scale indicates 1.0 substitutions per site for both phylogenies.

*Rps8* LRR-RK shares 55% (590/1,080) amino acid identity to Xa21 and is largely conserved in domain structure except for an additional LRR domain between seventh and eighth LRR domains of Xa21 and the presence of a 19 aa insertion in the kinase domain of *Rps8* LRR-RK (**Supplemental Figure 8**). The juxtamembrane domain is conserved in residues (S705, T707, and S708) that are required for proteolytic cleavage of Xa21 and interaction with Xb15 (S716), a protein phosphatase 2C (Park et al., 2008). *Rps8* LRR-RK has 20 predicted N-glycosylation sites (NxS/T) in the extracellular domain. Similar to Xa21, the *Rps8* LRR-RK is predicted to have serine-threonine specificity. The loss-of-function mutation *rps8-TM90* (G432R) is in a conserved residue (G414), whereas *rps8-TM2907* (A542T) is variable between *Rps8* LRR-RK (A542) and Xa21 (V542).

### *Rps8 Exo70* belongs to the Poales-specific *Exo70FX* subfamily

Exo70 is one of eight proteins which comprise the Exocyst complex along with Sec3, Sec5, Sec6, Sec8, Sec10, Sec15, and Exo84 (TerBush et al., 1996; Guo et al., 1999). The primary role of the Exocyst complex is tethering secretory vesicles to the plasma membrane in concert with soluble N-ethylmaleimide-sensitive factor attachment protein receptor (SNARE) proteins (Heider and Munson, 2012). Exo70 in vascular plants are highly expanded, spanning eleven clades (A, B, C, D, E, F, G, H, I, J, FX) (Synek et al., 2006; Cvrčková et al., 2012; Chi et al., 2015). In angiosperms, the majority of clades are conserved between monocot and dicot plants, although lineage-specific expansions have been observed for Exo70F in monocots and Exo70H in dicots. In contrast, Exo70FX is unique to monocots and exhibit substantial sequence divergence. To determine the phylogenetic relationship of the *Exo70* at *Rps8*, we identified, aligned, and constructed a maximum likelihood phylogenetic tree using a structure-guided approach for Exo70 proteins from Poaceae species with high quality reference genomes including *Hordeum vulgare* (barley), *Triticum aestivum* (wheat), *Brachypodium distachyon, Oryza sativa* (rice), *Setaria italica, Sorghum bicolor, Oropetium thomaeum*, and *Zea mays* (maize) (**Figure 7a**). Clades were annotated based on the previous classification for *O. sativa* and *B. distachyon* (Cvrčková et al., 2012). For all clades but Exo70FX, the original nomenclature was preserved and proteins from other species were annotated based on their relationships to existing clades and subclades (**Supplemental Table 7**). For the Exo70FX clade, bootstrap support varied substantially. To classify relationships within the Exo70FX, we found that the phylogenetic tree topology corresponded with syntenous relationships within the grasses. We integrated synteny as a criterion for delineating Exo70FX subclades. In total, 14 Exo70FX subclades were identified. *Rps8* Exo70 was designated Exo70FX12. Exo70FX12 is a single copy subclade located on chromosome 4H. Exo70FX12 is embedded within the Exo70FX11 subclade located on chromosome 2H (**Figure 7b; Supplemental Table 8**). Further analysis of the Exo70FX12 subclade found a second novel Exo70FX in barley that is absent in the reference genome but present in the barley accession Barke and located at a distinct locus on chromosome 2H. As this gene appears to be a novel transposition of the gene family, we designated it Exo70FX15. Collectively, these results show that the Exo70 at *Rps8* belongs to the Exo70FX clade, which is experiencing substantial intraspecific and interspecific expansion.

**Figure 7.**
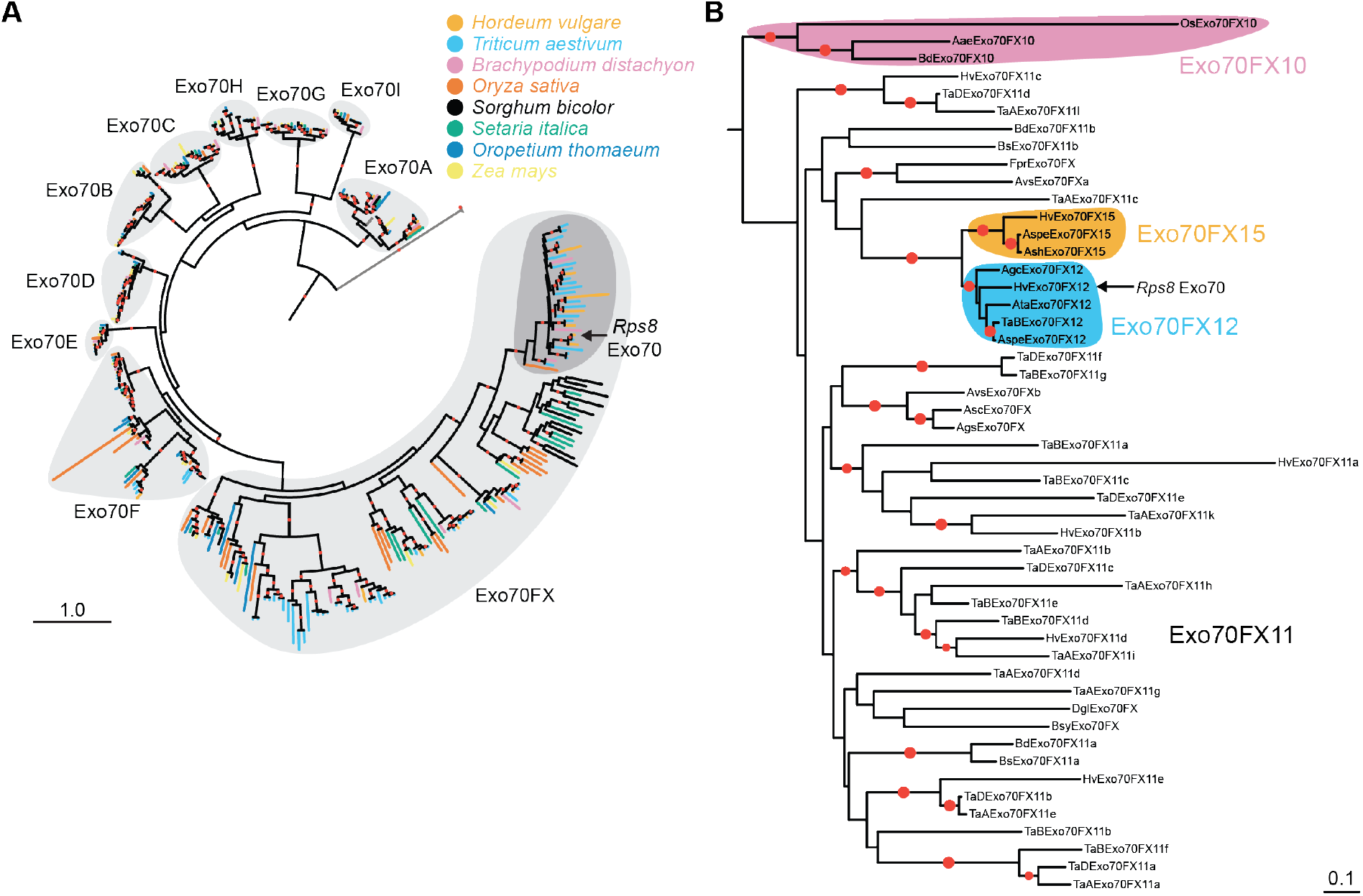
*Rps8* Exo70 belongs to the Poales-specific Exo70FX subfamily. **(A)** Maximum likelihood phylogenetic tree based on 365 full length Exo70 proteins from barley (*Hordeum vulgare*), wheat (*Triticum aestivum*), purple false brome (*Brachypodium distachyon*), rice (*Oryza sativa*), *Oropetium thomaeum*, maize (*Zea mays*), foxtail millet (*Setaria italica*), and sorghum (*Sorghum bicolor*). Structure-guided multiple sequence alignment was performed using MAFFT DASH with yeast, human, mouse, and *Arabidopsis thaliana* Exo70A1 proteins (**Supplemental Table 7**). The phylogenetic tree was rooted using yeast, human, and mouse Exo70 as outgroups. The ten Exo70 subfamilies (A, B, C, D, E, F, G, H, I, and FX) are shown in light grey regions and the Exo70FX10/11/12/15 clades is shown in dark grey. *Rps8 Exo70* encodes HvExo70FX12. Red dots indicate bootstrap support greater than 80%. Scale indicates 1.0 substitutions per site. **(B)** Maximum likelihood phylogenetic tree based on 51 full length Exo70 proteins from the Exo70FX10/11/12/15 clades and putative Exo70FX homologs from 13 additional Pooideae species (**Supplemental Table 8**). Color coding on the subtree indicates Exo70FX10 (pink), Exo70FX12 (blue), and Exo70FX15 (orange). The phylogenetic tree was rooted using the Exo70FX10 clade as an outgroup. Red dots indicate bootstrap support greater than 80%. Scale indicates 0.1 substitutions per site.

Previous work has shown that the Exo70FX clade is present in all evaluated Poaceae (grasses) species (Cvrčková et al., 2012). Our initial analysis encompassed genomes from eight Poaceae species. To ascertain the origin of the Exo70FX clade, we identified Exo70 clade members in the recently sequenced genomes of *Streptochaeta angustifolia* (Seetharam et al., 2021) and *Pharus latifolius* (Ma et al., 2021), which represent critical species at the boundary of the Poaceae family. In addition, we included *Ananas comosus* (pineapple; Poales, Bromeliaceae) and *Musa acuminata* (banana; Zingiberales, Musaceae) (D’Hont et al., 2012; Ming et al., 2015). Using the phylogenetic approach described above, integrating Exo70 from *H. vulgare, O. sativa, S. italica*, and *Z. mays*, we found conservation in all non-Exo70FX clades (**Supplemental Table 9**). In contrast, while the Exo70FX clade was present in *P. latifolius* and *S. angustifolia*, it was absent in *A. comosus* and *M. acuminata*. Therefore, the Exo70FX clade emerged after the divergence of the most recent ancestor of the Bromeliaceae-Poaceae.

The substantial diversification on Exo70FX prompted an examination of the conservation of subdomains within this family. In yeast, crystallization of Exo70 and Cryo-EM of the Exocyst complex has found that Exo70 is composed on five subdomains: CorEx, CAT-A, CAT-B, CAT-C, and CAT-D (Dong et al., 2005; Hamburger et al., 2006; Mei et al., 2018). In this model, the CorEx subdomain is intertwined with the CorEx subdomain of Exo84 to create the first-level heterodimer, which then forms a four-helix bundle with the CorEx subdomains of Sec10 and Sec15 (Mei et al., 2018). CAT-C and CAT-D of Exo70 interface with CAT-B and CAT-C of Sec5. Interaction with the plasma membrane via phospholipid binding is mediated by CAT-D (He et al., 2007; Liu et al., 2007). To assess variation in subdomain composition among Exo70, we compared the presence or absence of subdomains relative to *S. cerevisiae* Exo70. Using the amino acids coverage of Exo70 clades within the multiple sequence alignment, we found that there is family-specific variation in the presence/absence of CorEx and CAT-A sub-domains (**Figure 8; Supplemental Figure 9**). Exo70 clades Exo70A, Exo70B, Exo70C, Exo70F, Exo70G, and Exo70I show retention of all five sub-domains, whereas Exo70D, Exo70E, and Exo70H have a loss of coverage that suggests loss of the CorEx sub-domain. The Exo70FX clade has a unique coverage pattern, indicating that the majority of members have lost CorEx and CAT-A sub-domains. This observation was supported by a reduction in protein length in the majority of members in the Exo70FX clade (**Supplemental Figure 10**). Subdomain analysis of Exo70FX12 found that it lacked the canonical CorEx subdomain and likely has a truncated CAT-A subdomain. In summary, these results show that the Exo70FX clade is experiencing substantial species-specific expansion, high sequence divergence as compared to non-Exo70FX clades, and loss of the CorEx and CAT-A subdomain.

**Figure 8.**
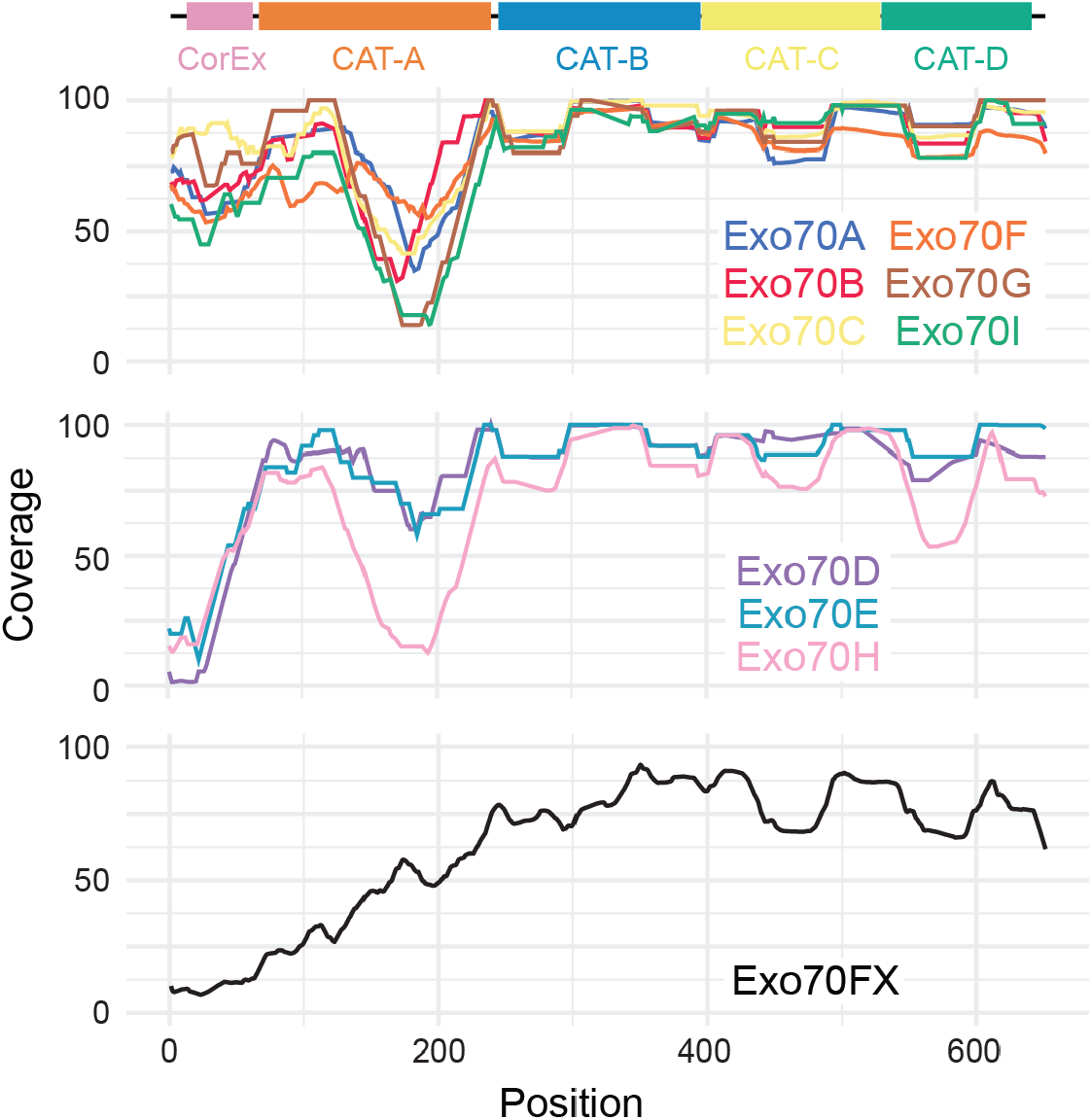
The majority of Exo70FX family members have lost several N-terminal subdomains. Coverage was estimated for each Exo70 clades using a sliding window of 50 amino acids based on the multiple sequence alignment used to generate the phylogenetic tree shown in Figure 7a. The x-axis is the position within the alignment and the y-axis is the percent coverage. Exo70 clades Exo70A (N=35), Exo70B (N=25), Exo70C (N=22), Exo70F (N=54), Exo70G (N=20), and Exo70I (N=10) show retention of all five sub-domains, whereas Exo70D (N=25), Exo70E (N=10), and Exo70H (N=14) have lost the CorEx sub-domain. The Exo70FX clade (N=143) has a unique coverage pattern with loss of the CorEx and CAT-A sub-domains in the majority of members. Sub-domain structure is based on yeast (Mei et al., 2018).

The diversity of sequence and structural information for Exo70 proteins enabled an evaluation of loss of function mutations relative to the subdomains of Exo70FX12. The mutations were found in the CAT-B (L130F; *rps8-TM3535*) and CAT-C (E271K; *rps8*.*b*) subdomains (**Figure 4e**). The position L130 is highly conserved in Exo70 proteins (297 of 352 conserved sites) including yeast and mouse, whereas E271 is the most common residue at this position in plant Exo70 proteins (102 of 379 conserved sites). Investigation of the predicted secondary structure based on yeast Exo70 and *A. thaliana* Exo70A1 found that both mutations occur in the C-terminal regions of alpha-helix domains. Thus, both loss-of-function mutations likely impact the secondary structure of Exo70FX12.

### *Exo70FX12* emerged in the Triticeae and exists as a *trans*-species presence/absence variation

*Rps8 LRR-RK* and *Exo70FX12* are located in a 546 kb region that exists as a presence/absence variation in domesticated (*H. vulgare* subsp. *vulgare*) and wild barley (*H. vulgare* subsp. *Spontaneum*). Our next analysis focused on addressing two questions: what is the evolutionary origin of *Exo70FX12* and does the locus exist in presence/absence variation in other species? To delineate the evolutionary origin of *Exo70FX12*, we searched sequenced genomes and *de novo* assembled transcriptomes of publicly available RNAseq data from the Bambusoideae, Oryzoideae, and Pooideae. Phylogenetic analysis with putative non-redundant homologs found *Exo70FX12* in species within the genera *Aegilops, Triticum, Agropyron*, and *Hordeum*, indicating that transposition of this gene from the *Exo70FX11* locus occurred in a common ancestor of the Triticeae (**Figure 7b**). A similar observation was made for *Exo70FX15*, which was found in the genera *Aegilops, Triticum*, and *Hordeum. Exo70FX12* was found in all sequenced *Triticum* B subgenomes (N=19), *Aegilops speltoides* (2 accessions; N=3), *Aegilops sharonensis* (2 accessions; N=16), *Agropyron cristatum* (1 accession; N=2), and *Aegilops tauschii* (4 accessions; N=16). *Rps8 LRR-RK* is found to be present with *Exo70FX12* in the majority of evaluated accessions. Based on these results, we conclude that *Rps8 LRR-RK* is ancient, with an origin prior to diversification of the BOP clade, whereas *Exo70FX12* is a recent innovation within the Triticeae.

## Discussion

We previously identified three *R* gene loci designated *Rps6, Rps7*, and *Rps8*, which provide resistance to the non-adapted pathogen *Pst* across a variety of cultivated barley accessions (Dawson et al., 2016; Li et al., 2016; Bettgenhaeuser et al., 2021). Fine-mapping of *Rps8* to a 936 kb locus on chromosome 4H identified a presence/absence variation across diverse barley accessions. Using a forward genetic screen, natural variation, and transgenic complementation, we found that *Rps8*-mediated resistance is conferred by a genetic module: *LRR-RK* and *Exo70FX12*, which are together necessary and required for *Rps8*-mediated resistance. *LRR-RK* belongs to the RK/Pelle LRR-XII subfamily and phylogenetic analysis of grasses found that it is the barley ortholog of rice *Xa21. Exo70FX12* belongs to the Exo70FX clade that emerged after the divergence of the most recent ancestor of the Bromeliaceae-Poaceae. *Exo70FX12* is a single copy Triticeae-specific locus with an origin through transposition of a member of the *Exo70FX11* locus, which itself contains multiple copies and experiences ectopic duplications. In contrast to other Exo70 clades, the Exo70FX clade exhibits several features such as loss of N-terminal CorEx and CAT-A subdomains, substantial sequence divergence among family members, and intraspecific presence/absence variation.

In green plants, transmembrane receptors and plant-specific kinases have experienced a massive expansion (Lespinet et al., 2002). Among these receptors, LRR-RK proteins are highly expanded and the predominant class of extracellular recognition receptor (Morillo and Tax, 2006; Fischer et al., 2016). The emergence of this gene family is associated with the colonization of land, an environment that would have exposed plants to a diverse array of microbial species (Lehti-Shiu et al., 2009). LRR-RK proteins are involved in regulating growth, abiotic stress responses, and extracellular recognition of pathogens (Shiu et al., 2004; Morillo and Tax, 2006; Tang et al., 2010; Fischer et al., 2016) and can be involved in multiple signaling pathways that facilitate crosstalk between growth and defense pathways, exemplified by the dual roles of the BAK1 co-receptor in immune and brassinosteroid signaling (Nam and Li, 2002; He et al., 2007). The LRR-RK gene family experiences species-specific and subclade-specific selective pressure, varying between purifying and expansion/diversifying, in a manner consistent with subclade-specific roles in growth and stress responses, as well as immunity (Lehti-Shiu et al., 2012; Fischer et al., 2016; Liu et al., 2017). The LRR-XII subfamily is associated with defense, and several well characterized LRR-RK genes involved in immunity, including FLS2, EFR and Xa21. Phylogenetic analysis of the RK Pelle LRR-XII subfamily found that *Rps8 LRR-RK* and *Xa21* are orthologs, sharing a common ancestor prior to the radiation of the Bambusoideae, Oryzoideae, and Pooideae (43 MYA, CI: 27 to 58 MYA (Kumar et al., 2017)). Expression of the wheat ortholog of *Rps8* LRR-RK, *TaXa21*, was found to be induced after heat treatment and transient virus-mediated silencing conferred a quantitative increase in *Pst* uredinia production in a temperature-dependent manner (Wang et al., 2019). Therefore, the recognition of *Pst* is likely an ancestral state of *Xa21* prior to the radiation of the Triticeae and implicates the *Xa21* orthogroup in resistance to adapted and non-adapted pathogens of wheat and barley, respectively. Among the RK LRR-XII subfamily in plants, *Rps8* LRR-RK is the first functionally validated family member to confer resistance to a fungal pathogen. FLS2 and EFR recognize well-conserved epitopes found in many bacteria, flg22 and elf18, respectively (Gomez-Gomez and Boller, 2000; Zipfel et al., 2004), whereas Xa21 recognizes a tyrosine-sulfated peptide derived from RaxX in *Xanthomonas oryzae* (Pruitt et al., 2015). While the ligand recognized by *Rps8* LRR-RK is unknown, this work supports the previously proposed hypothesis that the large expansion of LRR-XII genes in monocots is a shift towards recognizing ligands which are specific to a particular pathogen, in contrast with the characterized LRR-XII subfamily members of dicots (Shiu et al., 2004; Morillo and Tax, 2006).

The octomeric exocyst complex is found in fungi, metazoa, and plants with a primary role in localization and tethering of secretory vesicles to the cell membrane (TerBush et al., 1996; Guo et al., 1997; Kee et al., 1997). Genetic, functional, and structural analysis in yeast established that the exocyst complex is composed of eight proteins: Sec3, Sec5, Sec6, Sec8, Sec10, Sec15, Exo70, and Exo84 (TerBush et al., 1996; Guo et al., 1999). Within the complex, Exo70 interacts with Exo84, Sec10 and Sec15 via an N-terminal CorEx domain and with Sec5 via C-terminal CAT-C and CAT-D domains (Mei et al., 2018). Exo70 is an important hub for the exocyst complex, interacting with the plasma membrane (PI(4,5)P_2_) (He et al., 2007), SNARE proteins (Xu et al., 2013), RHO GTPase proteins (Roumanie et al., 2005), RAB GTPase proteins (Robinson et al., 1999; Koumandou et al., 2007), and Arp2/3 (Zuo et al., 2006) via C-terminal domains. Fusion with the membrane and secretion itself involves a variety of additional proteins, primarily components of the SNARE-complex (Sivaram et al., 2005; Yue et al., 2017). While animal and fungal genomes encode a single copy of the Exo70 gene, plants have evolved numerous copies which can be further divided into eleven clades: Exo70A, Exo70B, Exo70C, Exo70D, Exo70E, Exo70F, Exo70G, Exo70H, Exo70I, Exo70J, and Exo70FX (Synek et al., 2006; Cvrčková et al., 2012; Chi et al., 2015). These clades are non-redundant and exhibit clear evidence of subfunctionalization: they are expressed in different tissue types, localize to different membrane domains, and carry different cargoes (Li et al., 2010; Žárský et al., 2013; Žárský et al., 2019). While clades Exo70A, Exo70B, Exo70C, Exo70D, Exo70E, and Exo70G are well-conserved, the remaining clades exhibit variation between species. Exo70I, which is required for the establishment of mycorrhizal symbioses is not present in species which do not form these associations (Zhang et al., 2015). Exo70J is unique to legume species (Chi et al., 2015). Exo70H, which is involved in trichome development and defense is expanded in dicots (6-8 copies) relative to monocots (1-3 copies) (Cvrčková et al., 2012; Kulich et al., 2013; Kulich et al., 2015; Kubátová et al., 2019). Exo70F, conversely is expanded in monocots, and Exo70FX is unique to monocots (Cvrčková et al., 2012). Using recently released genomes within the Poales, phylogenetic analysis of the Exo70FX clade found that the clade emerged after the divergence of the most recent common ancestor of Bromeliaceae-Poaceae. *Exo70FX12* identified at *Rps8* is a member of the Exo70FX clade. To date, only one member of the Exo70FX clade has been characterized, *Exo70FX11b*, which was found to contribute towards penetration resistance against powdery mildew when transiently silenced (Ostertag et al., 2013). In barley, the *Exo70FX11* locus is located on chromosome 2H and contains six family members that show evidence of subfunctionalization based on tissue-specific expression (**Supplemental Figure 11**). *Exo70FX12* is derived from the *Exo70FX11* locus, which translocated to chromosome 4H prior to the radiation of the Triticeae as orthologs were found in *Agropyron cristatum, Aegilops tauschii, Aegilops speltoides*, and *Triticum aestivum* B genome. Several lines of evidence indicate that the Exo70FX clade is experiencing a different evolutionary trajectory as compared to other Exo70 families. First, the Exo70FX family experiences substantial sequence divergence, observed by long branch lengths on the phylogenetic tree, as well as extensive interspecific copy number variation. Second, of the ten Exo70 clades in grasses, only Exo70D and Exo70FX experience variable loss of the N-terminal region. For Exo70D, this is restricted to the CorEx domain, whereas Exo70FX proteins routinely lack CorEx and/or CAT-A domains. Third, we identified intraspecific variation for novel members of the Exo70FX clade, including Exo70FX12 and Exo70FX15. This contrasts with other clades, where no evidence exists for intraspecific presence/absence variation. Our current analysis has been restricted to the Pooideae, although we hypothesize that gene family expansion mediated by translocation may be common within other grass lineages for the Exo70FX clade. Fourth, Exo70F and Exo70FX are the only two clades identified as integrated domains in NLRs (Bailey et al., 2018) suggesting a role in immunity and the target of pathogen effectors. To the latter hypothesis, the rice blast effector AVR-Pii directly interacts with OsExo70F2 and OsExo70F3, and the interaction with OsExo70F3 is required for resistance mediated by the rice CC-NB-LRR pair Pii-1 and Pii-2 (Fujisaki et al., 2015). Lastly, *Exo70FX12* is required for *Rps8*-mediated resistance. This role depends on the *Rps8* LRR-RK, suggesting that *Exo70FX12* has evolved a specialized role. The Exo70FX11 family members appear to be insufficient to complement this function, despite several members having leaf expression. Collectively, this suggests that the Exo70FX clade is experiencing intense evolutionary pressure that may have its origins in adaptive evolution.

Extensive forward and reverse genetic screens have been performed on FLS2, EFR, and Xa21 to identify genes regulating their maturation, translocation, signaling, and degradation (Couto and Zipfel, 2016). These screens have uncovered proteins involved at different stages in the secretory pathway including the endoplasmic reticulum, the Golgi apparatus, and the trans-Golgi network (Li et al., 2009; Lu et al., 2009; Nekrasov et al., 2009; Chen et al., 2010; Park et al., 2010; Park et al., 2013). The final stage of transport to the plasma membrane involves the tethering of vesicles to the membrane mediated by the exocyst complex, followed by the fusion of the secretory vesicle catalyzed by the SNARE complex (He and Guo, 2009). The role of the exocyst complex in membrane receptor translocation was uncovered through reverse genetics in *A. thaliana* (Pečenková et al., 2011; Stegmann et al., 2012; Stegmann et al., 2013). Initially, mutants in *AtEXO70B2* were found to have greater susceptibility to a range of pathogens (Pečenková et al., 2011; Stegmann et al., 2013), and later found to be compromised in early responses to the elicitors flg22, elf18, chitin, and Pep1 (Stegmann et al., 2012). *A. thaliana* EXO70B1 is also required for flg22-mediated early immune responses, with other elicitors not tested (Stegmann et al., 2012). Further work found that AtEXO70B1 and AtEXO70B2 are essential for proper FLS2 homeostasis and trafficking to the membrane (Wang et al., 2020). EXO70B1 and EXO70B2 were found to directly interact with FLS2, as well as hetero-oligomerize (Wang et al., 2020). This role in membrane receptor signaling is likely conserved in monocots, at least for chitin recognition, as mutants in *OsExo70B1* have increased susceptibility to rice blast (*Magnaporthe oryzae*) and OsExo70B1 interacts directly with the chitin receptor CERK1 (Hou et al., 2020). Like FLS2, *Rps8* LRR-RK requires Exo70FX12, a member of the Poales-specific Exo70FX clade. The specialized role of Exo70FX12 contrasts with the conserved requirement of Exo70B family members in maintaining diverse receptor kinase abundance in the plasma membrane (Hou et al., 2020; Wang et al., 2020). As the Exo70B clade is conserved in all angiosperms (Cvrčková et al., 2012) and expressed in all cell layers of barley leaves, this indicates that Exo70FX12 role in *Rps8* LRR-RK signaling is distinct. In contrast to other Exo70 clades, the Exo70FX clade exhibits considerable interspecific and intraspecific expansion, high rates of sequence divergence between orthologs, inter-chromosomal duplications, and loss of N-terminal subdomains, including Exo70FX12. This latter observation is critical, as the N-terminal long coiled coil (CorEx) subdomain of yeast Exo70 interacts with Exo84 to form an antiparallel zipper that facilitates integration into the octomeric exocyst complex (Mei et al., 2018).

In plants, immune receptors such as RKs and NLRs experience expansion and diversification, likely due to the selective pressure exerted by plant pathogens. For the Exo70 gene family, tissue-specific expression likely underscored the early expansion of the gene family in plants (Žárský et al., 2009; Li et al., 2010; Žárský et al., 2019). As compared to other Exo70 clades, the rapid expansion and diversity of the Exo70FX clade may indicate that further subfunctionalization and/or neofunctionalization has occurred. Given that only two *Exo70FX* have been characterized (Ostertag et al., 2013), and both are involved in defense, it seems likely that this expansion is connected to an overall role in immunity. While the molecular function of the Exo70FX12 is unclear, its requirement in LRR-RK signaling implicates several possible models. In the first model, Exo70FX12 localizes LRR-RK to an appropriate domain of the plasma membrane with the involvement of other members of the Exocyst complex. Exo70FX12 would retain its capacity to interact with the plasma membrane and exocyst complex members. For the second model, Exo70FX12 localizes LRR-RK to the plasma membrane in an unconventional (non-exocyst complex) manner. Our third model is that the Exo70FX12 is not involved with localization of LRR-RK, and alternatively, Exo70FX12 is involved in signal transduction, perhaps as a scaffold to facilitate the interaction with other proteins. Different Exo70 clades are known to participate in diverse biological processes such as autophagy (Kulich et al., 2013; Acheampong et al., 2020) and mycorrhization (Zhang et al., 2015), that may or may not depend on its ancestral role as an exocyst complex member. This work has established a genetic function for Exo70FX12 in receptor kinase-mediated immunity and suggests that the Exo70FX clade may have evolved a specialized role in plant immunity.

### Materials and Methods

#### Plant and fungal materials

Barley accessions used in this study are referenced in **Supplemental Table 2**. SxGP DH-21, SxGP DH-47, and SxGP DH-103 are individuals from the SusPtrit x Golden Promise doubled-haploid population, which was provided by Rients Niks (Wageningen University, Netherlands) (Yeo et al., 2014). All plants were subjected to single seed descent prior to experimentation. *Pst* isolates 08/21 and 16/035 were collected in 2008 and 2016, respectively, in the United Kingdom and maintained at the National Institute of Agricultural Botany (NIAB). *Pst* isolates were increased on susceptible wheat cultivars, collected, and stored at 6°C.

#### Pathogen assays

*Puccinia striiformis* f. sp. *tritici* (*Pst*) inoculations were carried out by sowing seeds in groups of eight seeds per family, with four families spaced equidistantly around the rim of each 1 L pot of John Innes peat-based compost. Plants were grown at 18°C day and 11°C night using a 16 h light and 8 h dark cycle in a controlled environment chamber at NIAB, with lighting provided by metal halide bulbs (Philips MASTER HPI-T Plus 400W/645 E40). Barley seedlings were inoculated at 12 days after sowing, where first leaves were fully expanded and the second leaf was just beginning to emerge. Urediniospores of *Pst* were suspended in talcum powder, at a 1:16 ratio of urediniospores to talcum powder based on weight. Compressed air was used to inoculate seedlings on a spinning platform. After inoculation, seedlings were placed in a sealed bag and stored at 8°C for 48 h to increase humidity for successful germination of urediniospores. Subsequently, plants were returned to the growth chamber for the optimal development of *Pst* and phenotyped at 14 days post inoculation. Plants were scored using a 9-point scale from 0 to 4, with increments of 0.5, for chlorosis (discoloration) and infection (pustule formation) (Dawson et al., 2015). The scale indicates the percentage of leaf area affected by the corresponding phenotype where a score of 0 indicates asymptomatic leaves, i.e. no chlorosis, browning or pustules, and a score of 4 indicates leaves showing the respective phenotype over 100% of the surface area.

#### Microscopic phenotyping

Fixation, staining, and quantification of *Pst* in barley leaves was carried out according to Dawson *et al*. (Dawson et al., 2015). Briefly, barley leaves were collected at 14 dpi, placed in 1.0 M KOH with surfactant (Silwet L-77) and incubated at 37°C overnight (12-16 h). Leaves were washed three times in 50 mM Tris at pH 7.5. Leaf tissue was incubated in 1.0 mL of WGA-FITC solution (20 μg/mL WGA-FITC in 50 mM Tris at pH 7.5) overnight, mounted, and observed under blue light excitation on a fluorescence microscope with a GFP filter. pCOL estimates the percent of leaf colonized and pPUST the percent of leaf harboring pustules. Phenotyping was performed by evaluating the leaf surface in equally sized, adjacent portions. Within each field of view (FOV), the colonization of *Pst* was estimated to be less than 15%, between 15 and 50% or greater than 50% of the FOV area and given scores of 0, 0.5, or 1, respectively. The final pCOL score was determined by averaging these scores based on the total number of FOVs evaluated and ranged from 0 to 100%. pPUST was evaluated in a similar manner, but for the clustering pattern of *Pst* pustules. A 5x objective with a FOV of 2.72 mm x 2.04 mm was used.

#### Genetic analysis

DNA was extracted from leaf tissue using a 96-well plate-based CTAB based method (Bettgenhaeuser et al., 2021). Genotyping was performed using KASP assays at the John Innes Centre Genotyping Facility (Norwich, UK). Genetic maps for the Manchuria x Heils Franken F_2_ and BC_1_ populations were developed using markers spanning the consensus genetic map of barley approximately every 10 cM (**Supplemental Table 10**). Genetic distances were computed using MapManager QTX (v20) (Manly et al., 2001) using the Kosambi function. Two-point linkage tests (plotRF; R v4.1.1 (Team, 2013) and R/qtl v1.4.8-1 (Broman et al., 2003)) were used to evaluate genetic map integrity. Composite interval mapping was performed using QTLcartographer v1.17j (Basten et al., 1999) using five background markers, step size of 1 cM, and forward-background selection of background markers. Significance thresholds at α at 0.05 were determined using 1,000 permutations with resampling of background markers (Lauter et al., 2008).

The *Rps8* recombination screen was performed with 4,608 SxGP DH-21 x SxGP DH-103 F_2_ using the markers K1_1398 and K1_1470, which identified 127 recombinants. Marker saturation in the *Rps8* region was performed using the genomes of Barke, Bowman, Morex and Haruna Nijo and RNAseq data from SusPtrit and Golden Promise. The *Rps8* region was delimited to the flanking markers K_4819 and K_079610_445 and located in the barley genome using NCBI BLAST+ v2.2.31 (Altschul et al., 1990). Informative markers were applied to all recombinants derived from the recombination screens. A minimum of sixteen individuals from F_2:3_ families were independently assessed using *Pst* isolate 16/035.

#### Long-range assembly of CI 16139 chromosome 4H

Chromosome flow sorting of CI 16139 chromosome 4H was performed using the methods described by Doležel et al. (Doležel et al., 2012) and chromosomal high molecular weight (HMW) DNA was prepared as described in Thind et al. (Thind et al., 2017) (**Supplemental Figure 12**). Chicago Dovetail sequencing of the chromosome (Putnam et al., 2016) was performed by Dovetail Genomics (Santa Cruz, CA, USA), with initial assembly in Meraculous (v2.0.3) (Chapman et al., 2017) and final scaffolding in HiRise (Putnam et al., 2016). Briefly, Chicago libraries were generated by *in vitro* chromatin reconstitution using 250 ng HMW DNA. After fixation with formaldehyde, chromatin was digested with *Mbo*I, biotinylated nucleotides were added to 5’ overhangs, and proximity ligation was performed. After reversal of crosslinking the DNA was treated to remove biotin and sheared to approximately 400 bp. A sequencing library was generated using NEBNext Ultra enzymes (New England Biolabs) and Illumina-compatible adapters and sequenced using Illumina HighSeq X to produce 229 million paired end reads, which gave 95x coverage for chromosome 4H. An initial assembly using Meraculous had a length of 521.7 Mb across 57,043 scaffolds. The HiRise assembly and scaffolding had a length of 527.17 Mb across 1,702 scaffolds.

#### Genomic analyses

A GATK variant calling pipeline was used to identify variants in the barley accession Heils Franken relative to the reference genome (accession Morex). Briefly, raw reads were trimmed with Trimmomatic (v0.39) using parameters ILLUMINACLIP:TruSeq3-PE.fa:2:30:10, LEADING:5, TRAILING:5, SLIDINGWINDOW:4:15, and MINLEN:36 (Bolger et al., 2014). Alignment of reads onto the barley genome was performed with bwa mem (0.7.12-r1039) using parameters ‘-T 0 -M -R’ (Li, 2013). GATK (v4.2.0.0) (Van der Auwera and O’Connor, 2020) was used to sort (default parameters), mark duplicates (default parameters), perform variant calling using HaplotypeCaller with parameters ‘--dont-use-soft-clipped-bases --standard-min-confidence-threshold-for-calling 20’, and perform variant filtering using VariantFiltration with parameters QUAL>40, DP>9, and QD>20.0, which were based on assessment of the data sets distributions. The impact of variants on protein encoding genes was assessed with snpEff (v5.0d) with default parameters (Cingolani et al., 2012). Comparison of genomic regions encompassing the *Rps8* locus were performed using dotplots (dottup EMBOSS suite (Rice et al., 2000)) in Geneious Prime (v2021.2.2) (https://www.geneious.com). Repetitive DNA was annotated using RepeatMasker (v4.1.2-p1) (https://www.repeatmasker.org) using Dfam with RBRM (v3.2) (Storer et al., 2021), TREP (v19) (Wicker et al., 2002), and rmblastn (v2.11.0+).

#### Transcriptomic analysis

RNA was extracted from the first and second leaves of 10-day old plants. Tissue was harvested, frozen in liquid nitrogen, and ground to a fine powder using a mortar and pestle with grinding sand at -80°C. Ground tissue was suspended in TRI reagent, allowed to incubate for 5 min at room temperature before centrifugation at 12,000 *x g* to pellet the lysate. The supernatant was recovered and mixed with chloroform. This mixture was incubated at room temperature for 15 min before the phases were separated by centrifugation and the lighter phase was recovered. Nucleic acids were precipitated from the lighter phase with isopropanol, pelleted via centrifugation, then washed with 75% ethanol and resuspended in water. After extraction, RNA was purified, and checked for quality as described in (Dawson et al., 2016). RNA libraries were constructed using Illumina TruSeq RNA library preparation (Illumina; RS-122-2001) and sequenced using 100 or 150 bp paired end reads (Novogene). Trimming reads was performed with Trimmomatic (v0.39) with the following parameters: ILLUMINACLIP:2:30:10 using TruSeq3 paired end adapters, LEADING:5, TRAILING:5, SLIDINGWINDOW:4:15, and MINLEN:36. Spliced alignment of RNAseq reads to a DNA template was performed using hisat2 v2.2.1 using default parameters (Kim et al., 2019). Alignment of RNAseq reads to a cDNA template was performed using bwa mem (0.7.12-r1039) using default parameters. To identify sequence variation in *Rps8 LRR-RK* and *Exo70FX12*, the QKgenome_conversion.py pipeline was used (Moscou, 2021). Briefly, reads were aligned to the 936 kb *Rps8* region of the sequenced Morex genome using hisat2 (v2.2.1) with the default parameters. samtools v1.11 (Li et al., 2009) was used to compress and sort reads. The number of reads mapping to each nucleotide position within the locus was calculated using bedtools genomecov (Quinlan and Hall, 2010). QKgenome_conversion.py integrates variant calling and coverage to identify variants in annotated genes with coverage greater than or equal to 10 reads. *De novo* transcriptomes were assembled using Trinity (v2.4.0) using default parameters (Grabherr et al., 2011). For tissue-specific expression, RNAseq data was processed using Trimmomatic, as described above, and kallisto (0.46.0) (Bray et al., 2016) using the barley predicted transcriptome (version 3) to determine expression level (transcripts per million) using 100 bootstraps (Mascher et al., 2021).

#### Construct development and plant transformation

*Exo70FX12* coding sequence and its native promoter and terminator were amplified from barley Golden Promise gDNA using Phusion High-Fidelity DNA Polymerase (NEB). PCR primers were designed 2 kb upstream of the predicted start of transcription, and 1.5 kb downstream of the predicted stop codon. PCR was performed by assembling a reaction mix containing 2.5 μL Phusion Master Mix (NEB), 2.5 μL dNTPs (200 μM), 1 μL forward and reverse primer (SH_12_p1f and SH_12_p1r; 400 nM each), 1 μL DNA (100 ng gDNA or 10 ng plasmid DNA), and 0.2 μL Phusion Taq polymerase (NEB) per reaction vessel using a BioRad G-Storm GS4 thermocycler, set to cycle through: 1.5 min at 94 °C, 35 repeats of 30 sec at 94 °C, 30 sec at 50°C, 30 sec per kb at 72 °C, 5 minutes at 72 °C, and a cooling stage of 10 °C. PCR product was gel excised and extracted prior to cloning into pCR-XL-2-TOPO vector (Invitrogen). Plasmids were extracted from *E. coli* transformed cells and sequenced using Sanger sequencing (Genewiz, Oxford, UK). The complete *Exo70FX12* transcriptional unit was then assembled into pBract202 binary vector (BRACT) using the Gibson method (Gibson et al., 2009). Briefly, fragments harbouring 20 bp overlapping overhangs were PCR amplified from the *Exo70FX12* transcriptional unit and the acceptor pBract202 vector, then purified and mixed in an equimolar ratio (200 ng of total DNA) with 15 μL home-made 1.33x one-step isothermal DNA assembly master mixture prepared as described in Gibson *et al*. (2009). Reaction was incubated at 50°C for one hour followed by transformation into *E. coli* heat-shock competent cells. Plasmids were extracted from *E. coli* transformed cells.

Constructs with *LRR-RK* and *Exo70FX12* + *LRR-RK* constructs were assembled using the Golden Gate cloning method (Engler et al., 2008). Domesticated (removal of internal *Bsa*I/*Bbs*I sites without changes in encoded amino acids) coding sequence, native promoter and native terminator DNA parts were synthesized (Twist Bioscience) and separately cloned into Golden Gate compatible level 0 vectors. Restriction-ligation reactions (15 μL) with 1x T4 DNA ligase buffer (NEB), 1.5 μg BSA, 5U *Bbs*I (Thermo Scientific), 1000U T4 DNA ligase (NEB), 100 ng acceptor vector and twice the molar amount of the DNA part. Reactions were performed for 26 cycles of alternating incubations at 37 °C for 3 min and 16 °C for 4 min, followed by 5 min at 50°C and 5 min at 80°C to inactivate the enzymatic components. After *E. coli* transformation, corresponding transcriptional units were assembled into Golden Gate level 1 vectors by similar restriction-ligation procedure but using *Bsa*I instead of *Bbs*I. The *Exo70FX12HF* + *LRR-RK* construct was generated from the *Exo70FX12* + *LRR-RK* construct by site-directed mutagenesis. Final constructs were subsequently assembled into pICSL4723 Golden Gate compatible binary vector (AddGene #48015) by *Bbs*I-T4 DNA ligase restriction-ligation reactions. All primers used for molecular cloning are described in **Supplemental Table 11**.

Assembled constructs were then introduced into *A. tumefaciens* strain AGL1 by electroporation. Barley plants transformation was performed using the *Agrobaterium*-mediated transformation method described by Hensel et al. (Hensel and Kumlehn, 2004). T-DNA insert copy number testing was performed by iDna Genetics (Norwich, UK) by quantitative real time PCR using the selectable marker gene *hyg* similar to the approach in (Bartlett et al., 2008).

#### Domain, subdomain, and phylogenetic analysis of Exo70

To identify proteins containing Exo70 domains, InterProScan v5.36-75.0 (Jones et al., 2014) using default parameters was used on predicted transcripts and annotated CDS. Proteins annotated with the Exo70 Pfam family (PF03081) were extracted. Structure-guided multiple sequence alignment of Exo70 proteins was performed using MAFFT (v7.481) (Rozewicki et al., 2019) using DASH with default parameters. Exo70 structures included in the alignment were derived from *Arabidopsis thaliana* Exo70A1 (PDB 4RL5) (Zhang et al., 2016), *Saccharomyces cerevisiae* (yeast) Exo70 (PDB 2B1E and 5YFP) (Dong et al., 2005; Mei et al., 2018), and *Mus musculus* (mouse) (PDB 2PFT) (Moore et al., 2007). The phylogenetic tree was constructed using RAxML (v8.2.12) (Stamatakis, 2014) with the JTT amino acid substitution model, gamma model of rate heterogeneity, and 1,000 bootstraps. A convergence test performed using RAxML autoMRE found convergence after 200 bootstraps. iTOL (Letunic and Bork, 2021) was used for phylogenetic tree visualization, and *S. cerevisiae* and *M. musculus* Exo70 were used as outgroups. For the subdomain analysis of Exo70, a domain was considered present in if at least 30 total residues were present over the alignment region corresponding to that domain in *S. cerevisiae* Exo70. Protein modelling was performed with Phyre2 using normal modelling mode (Kelley et al., 2015).

#### Phylogenetic analysis of RK LRR-XII subfamily

To identify RK LRR-XII subfamily in diverse grass species, we used the Hidden Markov Model (HMM)-based approach developed by Lehti-Shiu and Shiu (2012). Briefly, all kinase domain containing proteins were identified using 38 kinase Pfam identifiers (**Supplemental Table 12**). Grass species genomes were accessed from diverse sources (**Supplemental Table 4**). *De novo* assembled transcriptomes were generated for species with available leaf RNAseq data sets (**Supplemental Tables 2 and 5**). Putative kinases were classified with HMMs based on individual kinase families using hmmalign (HMMER 3.1b2) (Johnson et al., 2010). Multiple sequence alignment was performed using kalign with default parameters (Lassmann and Sonnhammer, 2005). The QKphylogeny_alignment_analysis.py script was used to filter the alignment for variable sites represented in at least 20% of proteins and sequences spanning at least 40% of the alignment length (https://github.com/matthewmoscou/QKphylogeny). Maximum likelihood phylogenetic tree construction was performed using RAxML using the JTT amino acid substitution model, gamma model of rate heterogeneity, and 1,000 bootstraps. Visualization of the phylogenetic tree was performed with iTOL.

## Acknowledgements

We greatly appreciated valuable discussions with Christine Faulkner, Sophien Kamoun, Ralph Hückelhoven, Viktor Žárský, Cristobal Uauy, Sebastian Schornack, Jack Rhodes, and Cyril Zipfel. Photography was supported by Andrew Davis and Phil Robinson. Assistance in the greenhouse was provided by Sue Banfield and the John Innes Centre Horticultural team. Seed was kindly provided by Rients Niks, Wendy Harwood, and the National Small Grains Collection (USDA-ARS). We thank P. Cápal, M. Said, Z. Dubská, J. Weiserová and E. Jahnová (Institute of Experimental Botany, Olomouc) for assistance with chromosome flow sorting and preparation of chromosome DNA. Funding for this research includes United Kingdom Research and Innovation-Biotechnology and Biological Sciences Research Council Norwich Research Park Doctoral Training Partnership (grant no. BB/M011216/1 to SH, BB/F017294/1 to JB) and Institute Strategic Programme (grant no. BB/P012574/1 to MJM and BBS/E/J/000PR9795 to MJM), Fulbright Commission Postgraduate Award (MB), Human Frontier Science Program Long-term Fellowship (grant no. LT000218/2011-L to MJM), European Regional Development Fund (grant no. CZ.02.1.01/0.0/0.0/16_019/0000827 to IM, JD and HŠ), and Gatsby Charitable Foundation (MJM).

## Data Availability

The RNAseq data generated in this study are found in the NCBI database under BioProject codes PRJNA292371, PRJNA376252, PRJNA378334, and PRJNA378723. The genome assembly and sequencing data for barley accession CI 16139 chromosome 4H generated in this study have been deposited in the NCBI database under BioProject code PRJNA786761. The whole genome shotgun sequencing data for barley accession Heils Franken generated in this study have been deposited in the NCBI database under BioProject code PRJNA787282. The sequences of plasmids used for plant transformation in this study have been deposited in the NCBI database with accession codes OL791272 (Exo70), OL791270 (LRR-RK), OL791269 (LRR-RK+Exo70FX12), and OL791271 (LRR-RK+Exo70FX12HF) and Figshare (Holden and Moscou, 2021). Genotypic, phenotypic, and source data for figures and supplemental figures have been deposited on Figshare (Holden and Moscou, 2021). A material transfer agreement with The Sainsbury Laboratory is required to receive the materials. The use of the materials will be limited to non-commercial research uses only. Please contact MJM regarding biological materials, and requests will be responded to within 60 days.

## Code availability

QKcartographer, QKgenome, and QKphylogeny suite of Python scripts are maintained on GitHub (https://github.com/matthewmoscou/QKcartographer; https://github.com/matthewmoscou/QKgenome; https://github.com/matthewmoscou/QKphylogeny) and Figshare (Moscou, 2021, 2021).

## Competing Interests

The authors declare no competing interests.

## Supplemental Tables

**Supplemental Table 1**. Protein encoding genes in the *Rps8* region

**Supplemental Table 2**. Association of barley diversity and *Rps8*-mediated resistance

**Supplemental Table 3**. Loss of function mutants in *Rps8*-mediated resistance

**Supplemental Table 4**. Genomes of monocot species used for gene family analysis

**Supplemental Table 5**. Presence/absence variation of *Rps8 LRR-RK* and *Exo70FX12* in diverse grass species

**Supplemental Table 6**. LRR-RK family XII receptor kinases from diverse grass species and their relationship to *Xa21*

**Supplemental Table 7**. Annotation of Exo70 families in eight Poaceae species

**Supplemental Table 8**. Exo70FX family members in the Exo70FX10, Exo70FX11, Exo70FX12, and Exo70FX15 clades in diverse grass species

**Supplemental Table 9**. Exo70 families in diverse Poales species

**Supplemental Table 10**. Genetic markers used to map and identify *Rps8*

**Supplemental Table 11**. Primers used for molecular cloning

**Supplemental Table 12**. Pfam identifiers for kinase domains

**Supplemental Figure 1.**
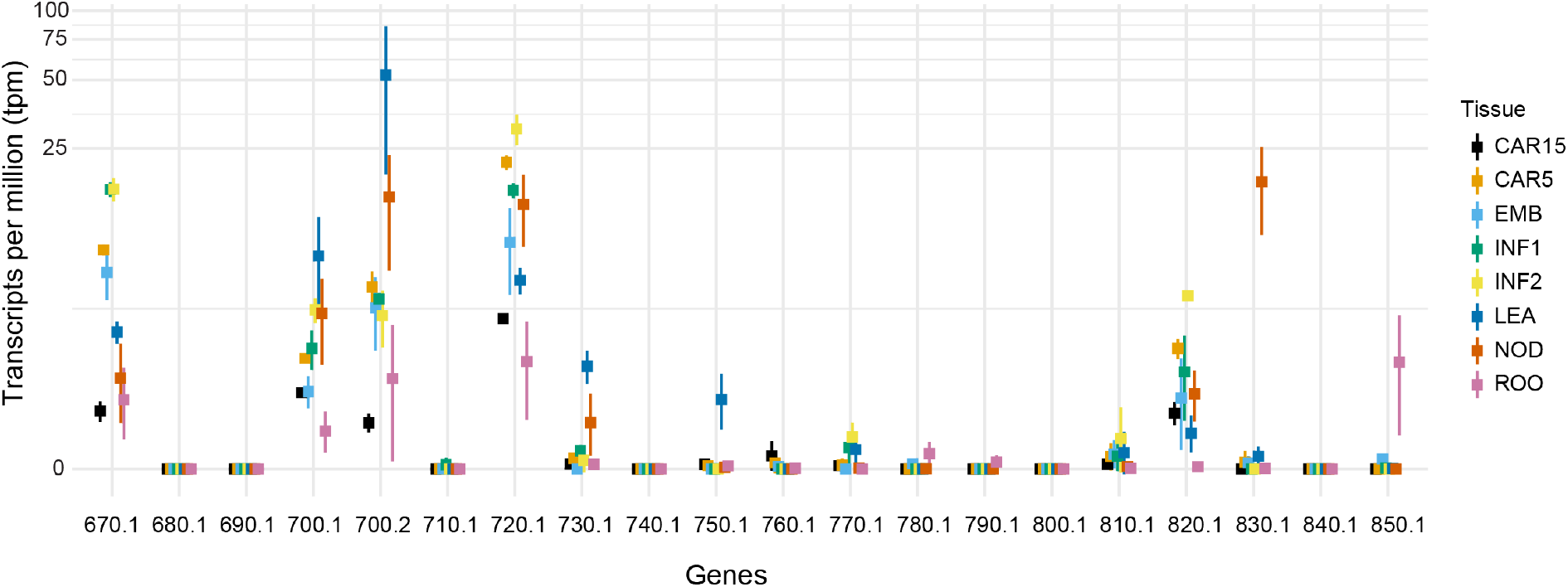
Expression analysis of genes in the *Rps8* locus in eight diverse tissues. RNAseq data was obtained from EMBL/ENA accession ERP001600 (IBGSC, 2012), trimmed, and used to estimate expression level (transcripts per million; y-axis; pseudo-log scale) based on the barley predicted transcriptome (Mascher et al., 2021). Identifiers (x-axis) correspond to the suffix of genes within the *Rps8* locus based on the longer identifier HORVU.MOREX.r3.4H0407xxx. Mean and standard deviation are shown as square and whiskers. Color coding and order shows the tissue including early developing grain (15 days post anthesis (CAR15) and 5 (CAR5)), germinating grain (4 day) embryos (EMB), early developing inflorescences (5 (INF1) & 15 mm (INF2)), shoots from seedlings (LEA; 10 cm stage), developing tiller internodes (NOD) (six-leaf stage), and roots from seedlings (ROO; 10 cm stage).

**Supplemental Figure 2.**
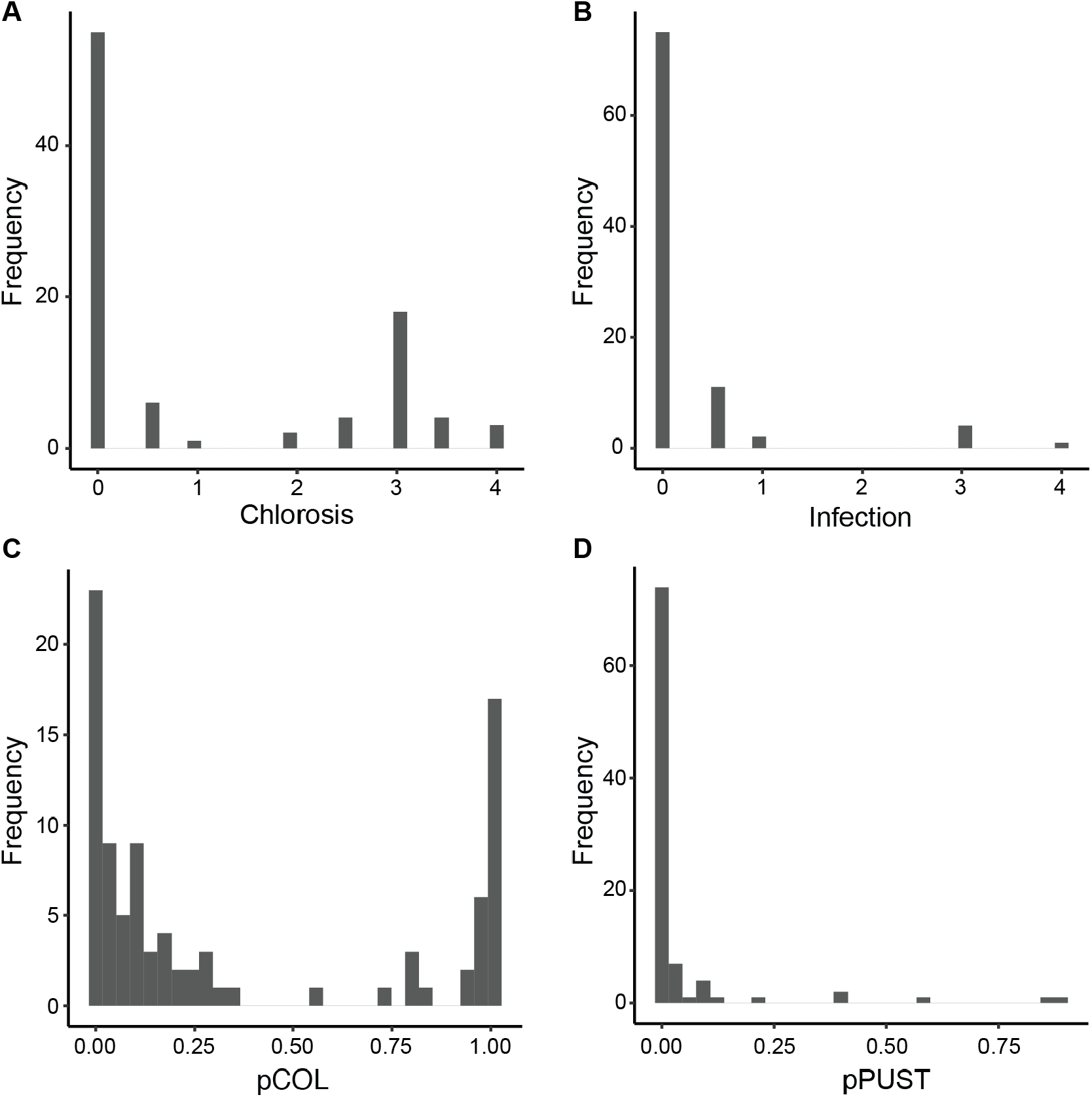
Phenotypic distributions of disease traits in the Manchuria x Heils Franken F_2_ population inoculated with *P. striiformis* f. sp. *tritici* isolate 08/21. Histograms showing the phenotypic distributions for macroscopic (A) chlorosis and (B) infection and microscopic colonization (pCOL) (C) and pustule formation (pPUST) (D). The population is composed of 94 F_2_ individuals.

**Supplemental Figure 3.**
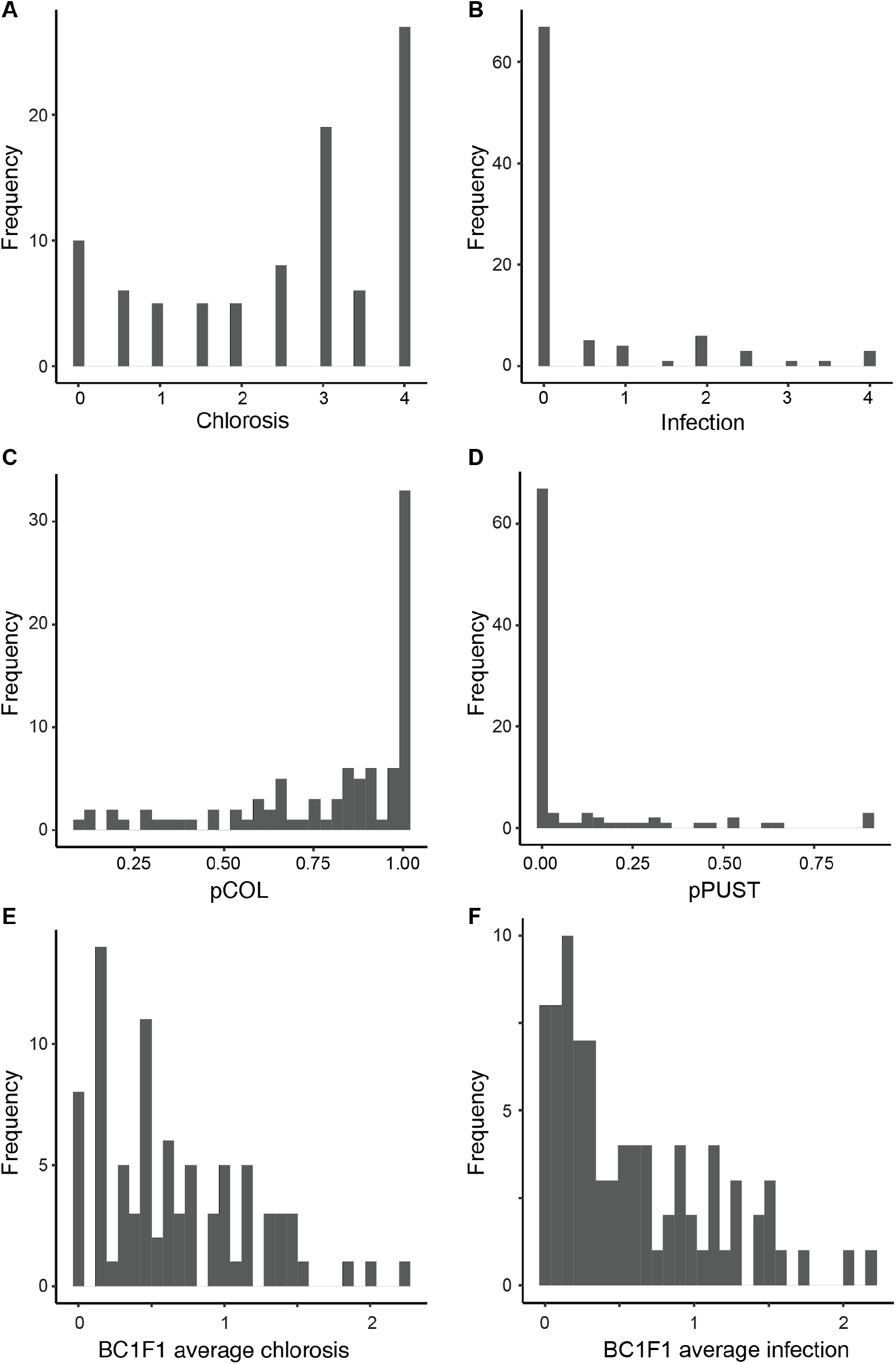
Phenotypic distributions of resistance traits in the Manchuria x Heils Franken BC_1_ population and BC_1_F_1_ progeny inoculated with *P. striiformis* f. sp. *tritici* isolate 08/21. Histograms showing the phenotypic distributions for macroscopic (A) chlorosis and (B) infection and microscopic colonization (pCOL) (C) and pustule formation (pPUST) (D) of the Manchuria x Heils Franken BC_1_ population (N=94), and average chlorosis (E) and infection (F) phenotypes for the Manchuria x Heils Franken BC_1_F_1_ population (N=85; 8 seedlings per family).

**Supplemental Figure 4.**
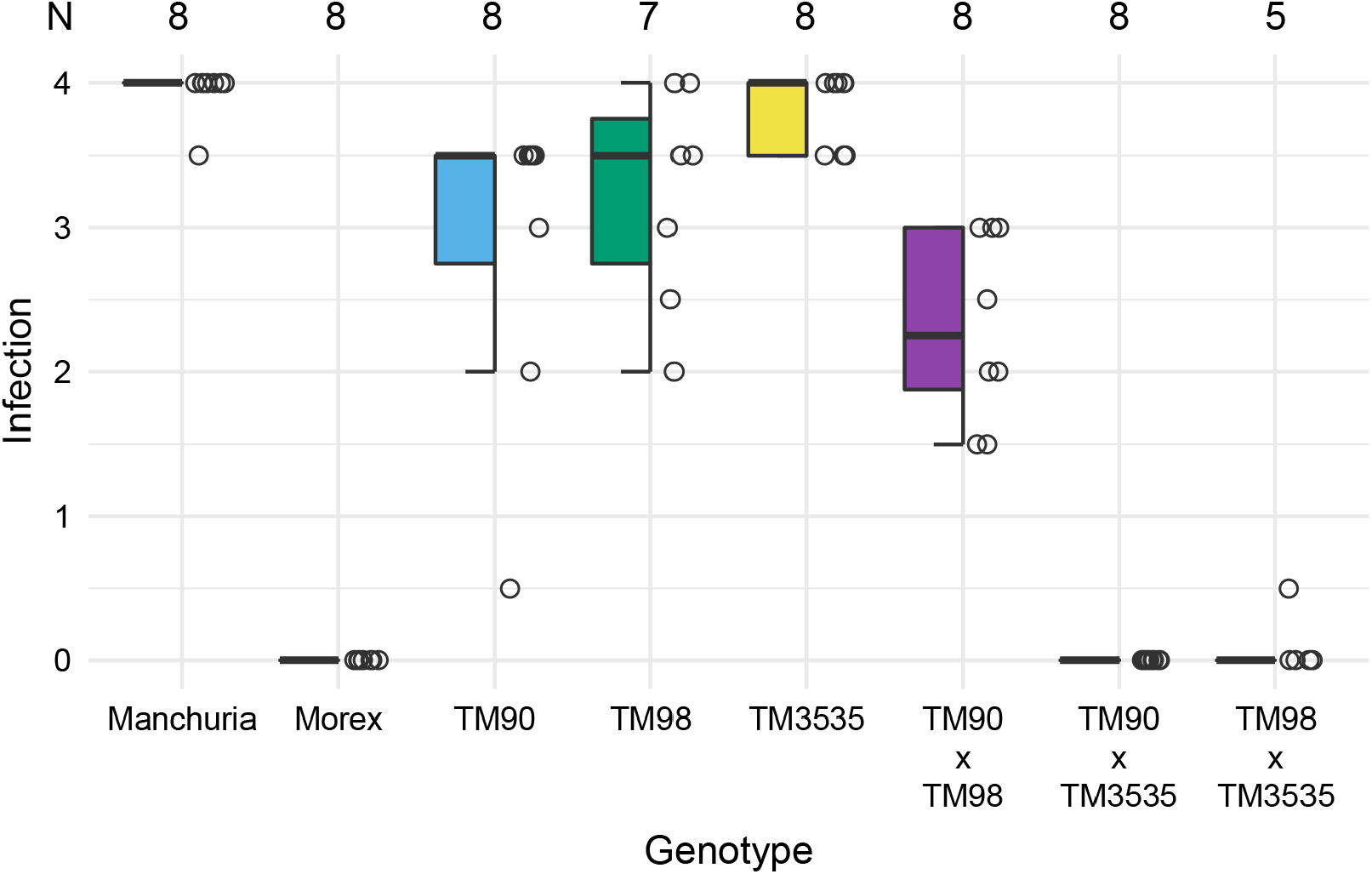
Complementation tests for *rps8* mutants TM90, TM98, and TM3535. Infection phenotypes (y-axis) of confirmed mutants and pairwise F_1_ progeny inoculated with *Pst* isolate 16/035. For each genotype (x-axis), boxplot and individual data points are shown from a single experiment that included F_1_ individuals. Total number of evaluated individuals (N) is shown. Data for controls Manchuria, Morex, TM90, TM98, and TM3535 is a subset of data shown in Figure 4d.

**Supplemental Figure 5.**
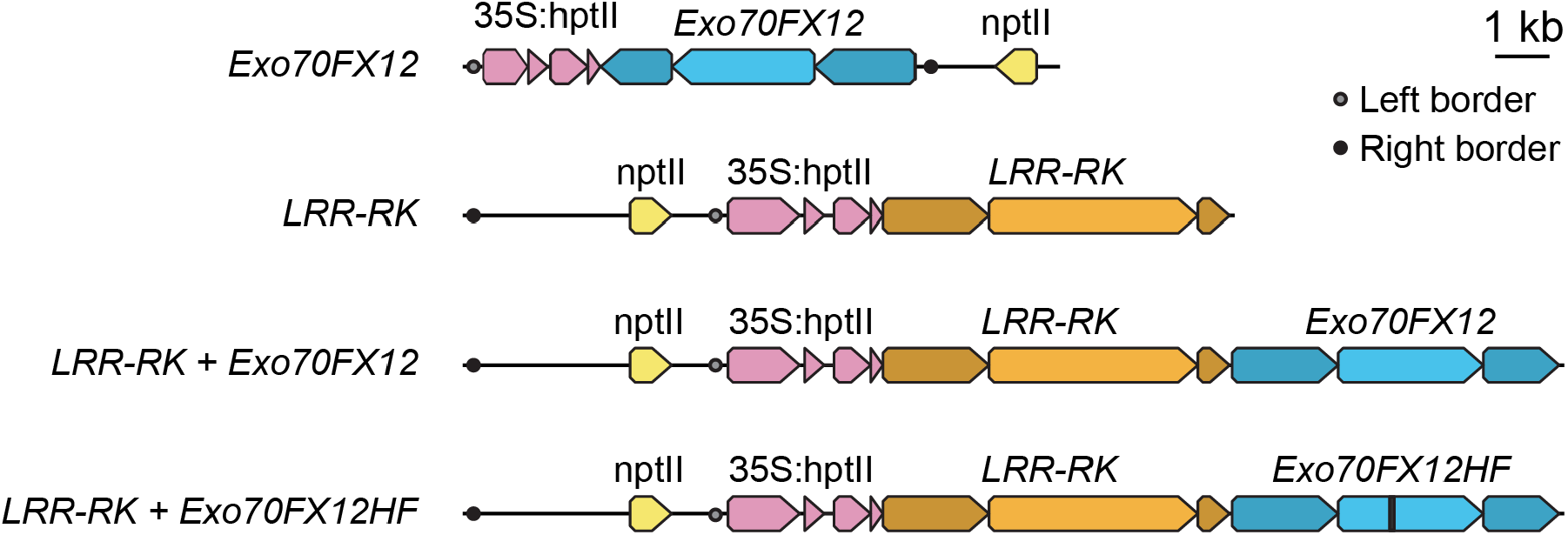
Transformation constructs for candidate genes at the *Rps8* locus. The *Exo70FX12* construct was generated using a PCR fragment encompassing the native genomic context of *Exo70FX12* with approximately 2 kb promoter and 1.5 kb terminator. The constructs *LRR-RK* and *LRR-RK* + *Exo70FX12* were generated by synthesizing domesticated fragments of native genomic context with approximately 2 kb promoter and 0.6 kb (*LRR-RK*)/1.5 kb (*Exo70FX12*) terminator and assembled using Golden Gate cloning. *LRR-RK* + *Exo70FX12HF* was generated using site-directed mutagenesis using the *LRR-RK* + *Exo70FX12* construct as template, recreating the Heils Franken allele for *Exo70FX12*. All constructs used nptII as a bacterial selectable marker (shown in yellow) and hptII driven by the 35S Cauliflower Mosaic Virus promoter for plant selection during transformation (shown in pink). Left and right T-DNA borders are shown with filled grey and black circles, respectively.

**Supplemental Figure 6.**
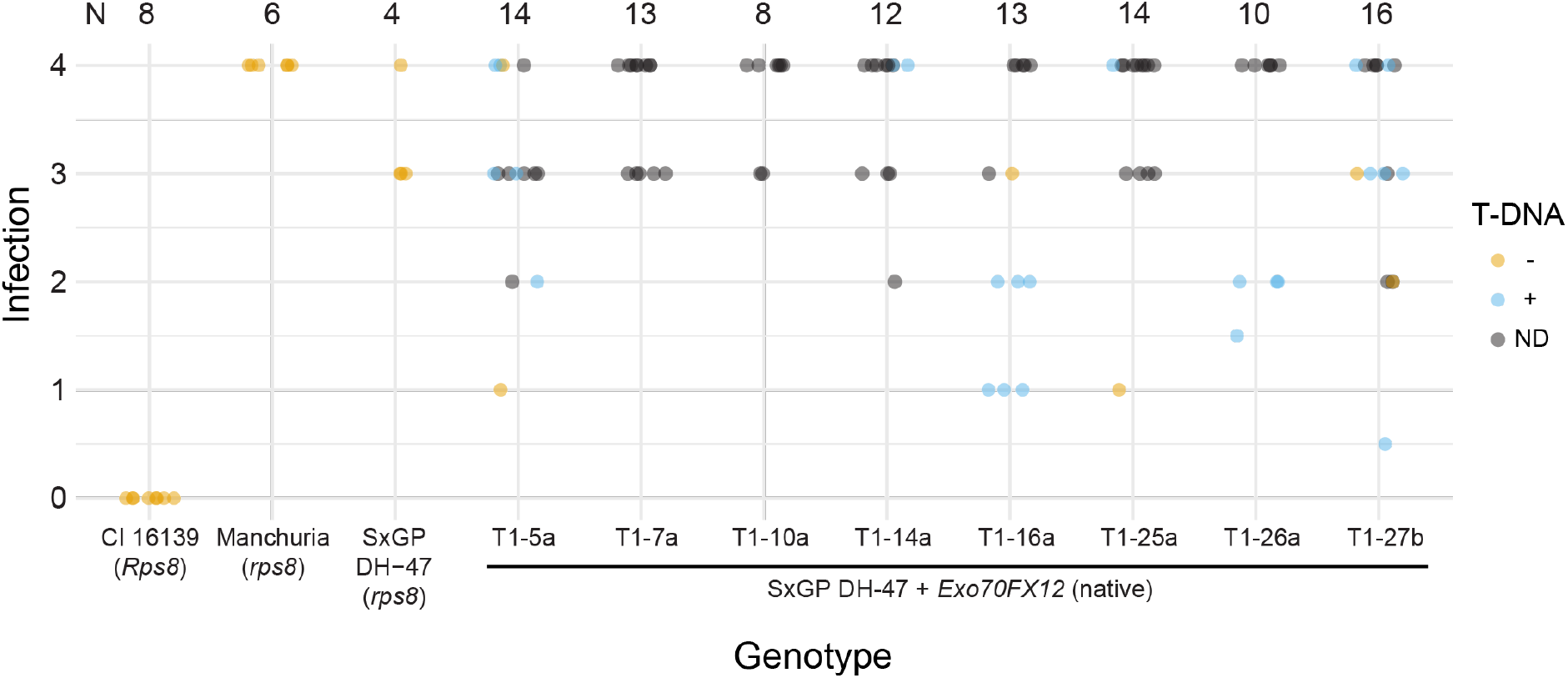
Infection phenotypes for T_1_ transgenic lines natively expressing *Exo70FX12*. Infection phenotypes for independent single copy transformants of barley accession SxGP DH-47 expressing *Exo70FX12* under native promoters and terminators. Presence (blue) or absence (orange) of the T-DNA was determined using a qPCR-based assay. When not determined (ND), data points are in black. Inoculations were performed using *Pst* isolate 16/035 and scored at 14 days post inoculation, N shows the number of evaluated seedlings, and each panel represents a single experiment.

**Supplemental Figure 7.**
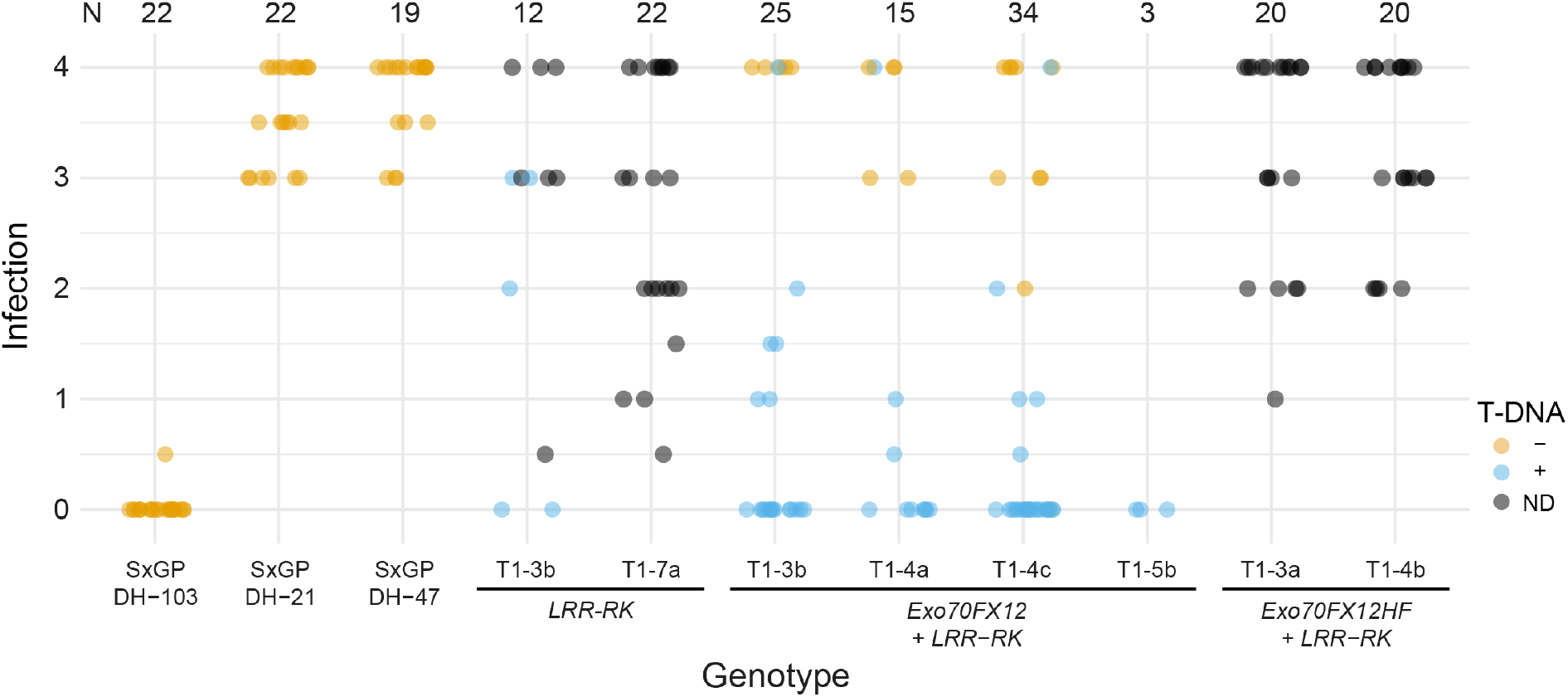
Extended set of transgenic lines expressing *LRR-RK, LRR-RK* and *Exo70FX12*, and *LRR-RK* with *Exo70FX12* allele from Heils Franken. Infection phenotypes for independent single copy transformants of barley accession SxGP DH-47 expressing *LRR-RK, Exo70FX12* and *LRR-RK*, and *Exo70FX12* (Heils Franken allele) and *LRR-RK*, under their native promoters and terminators. Presence (blue) or absence (orange) of the T-DNA was determined using a qPCR-based assay. When not determined (ND), data points are in black. For both panels, inoculations were performed using *Pst* isolate 16/035 and scored at 14 days post inoculation, N shows the number of evaluated seedlings, and each panel represents a single experiment.

**Supplemental Figure 8.**
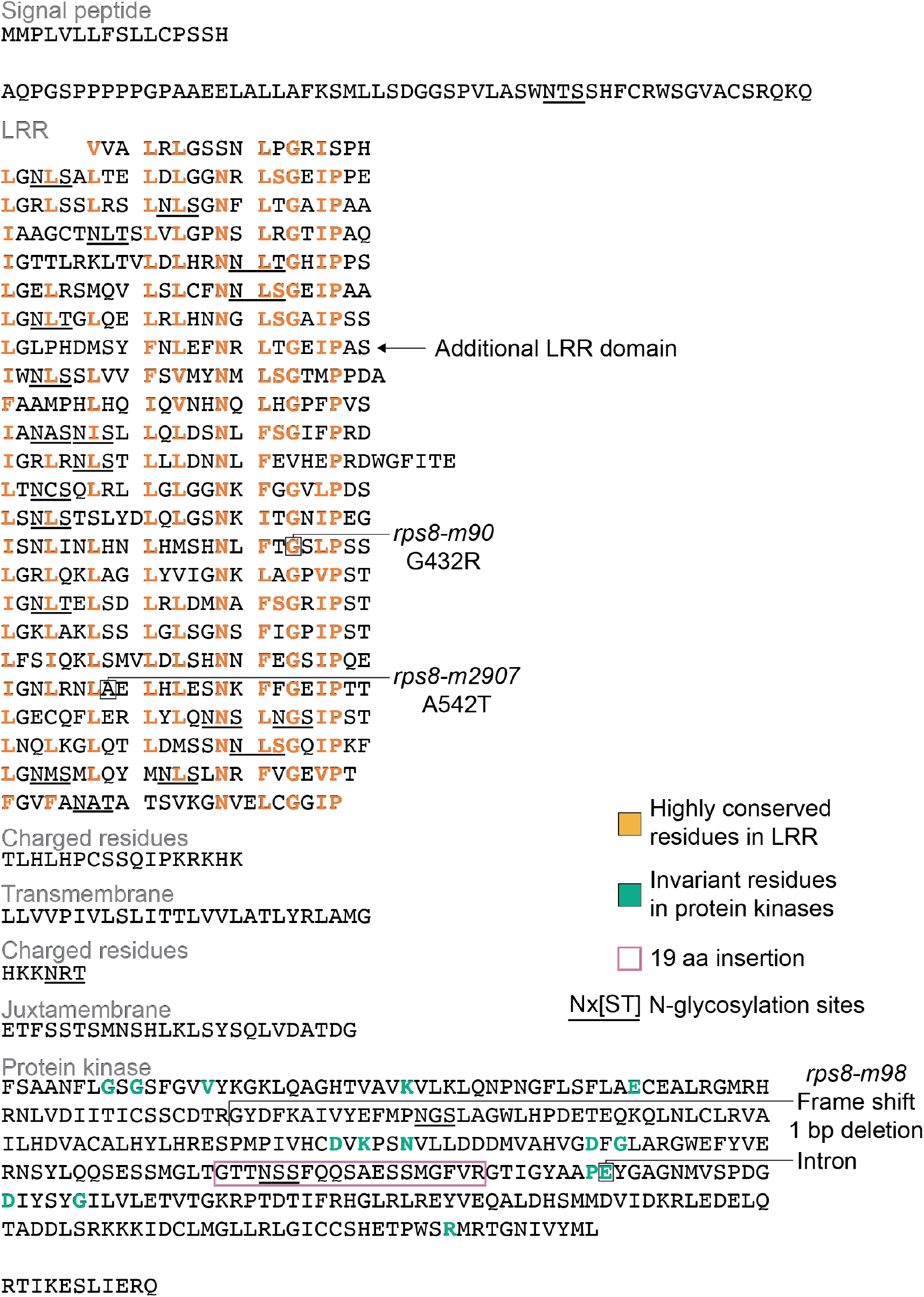
Domain analysis of *Rps8* LRR-RK. Domains of *Rps8* LRR-RK were annotated based on InterProScan and alignment with Xa21. The gene model used for *Rps8* LRR-RK is HORVU.MOREX.r3.4HG0407750.1. Highly conserved residues in the LRR are shown in bold orange, invariant residues in the protein kinase domain shown in bold green, the site of individual mutations (*rps8-TM90, rps8-TM98, rps8-TM2907*) and their result change to the protein, putative N-glycosylation sites, the position of the single intron, and the 19 amino acid insertion in the protein kinase domain in a pink rectangle. Design of the figure is based on the original characterization of Xa21 (Song et al., 1995).

**Supplemental Figure 9.**
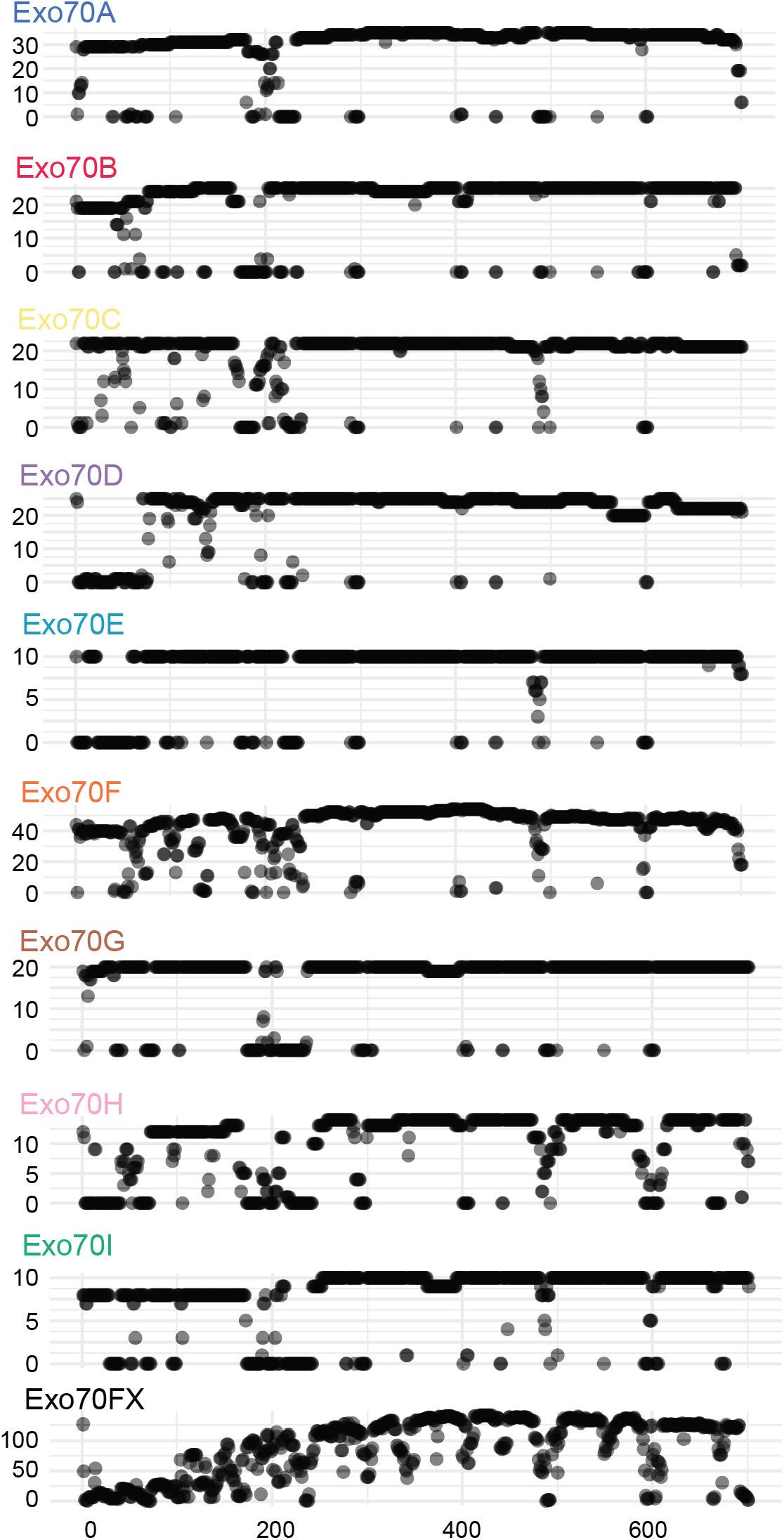
The Exo70FX family members have a distinct pattern of N-terminal loss as compared to other Exo70 families. Coverage was computed on every position in the multiple sequence alignment used to generate the phylogenetic tree shown in Figure 7a. The x-axis is the position within the alignment and the y-axis is the coverage based on the total number of family members. Exo70 families Exo70A (N=35), Exo70B (N=25), Exo70C (N=22), Exo70F (N=54), Exo70G (N=20), and Exo70I (N=10) show retention of all five sub-domains, whereas Exo70D (N=25), Exo70E (N=10), and Exo70H (N=14) have lost the CorEx sub-domain. The Exo70FX family (N=143) has a unique coverage pattern with loss of the CorEx and CAT-A sub-domains in the majority of members. Sub-domain structure is based on yeast (Mei et al., 2018).

**Supplemental Figure 10.**
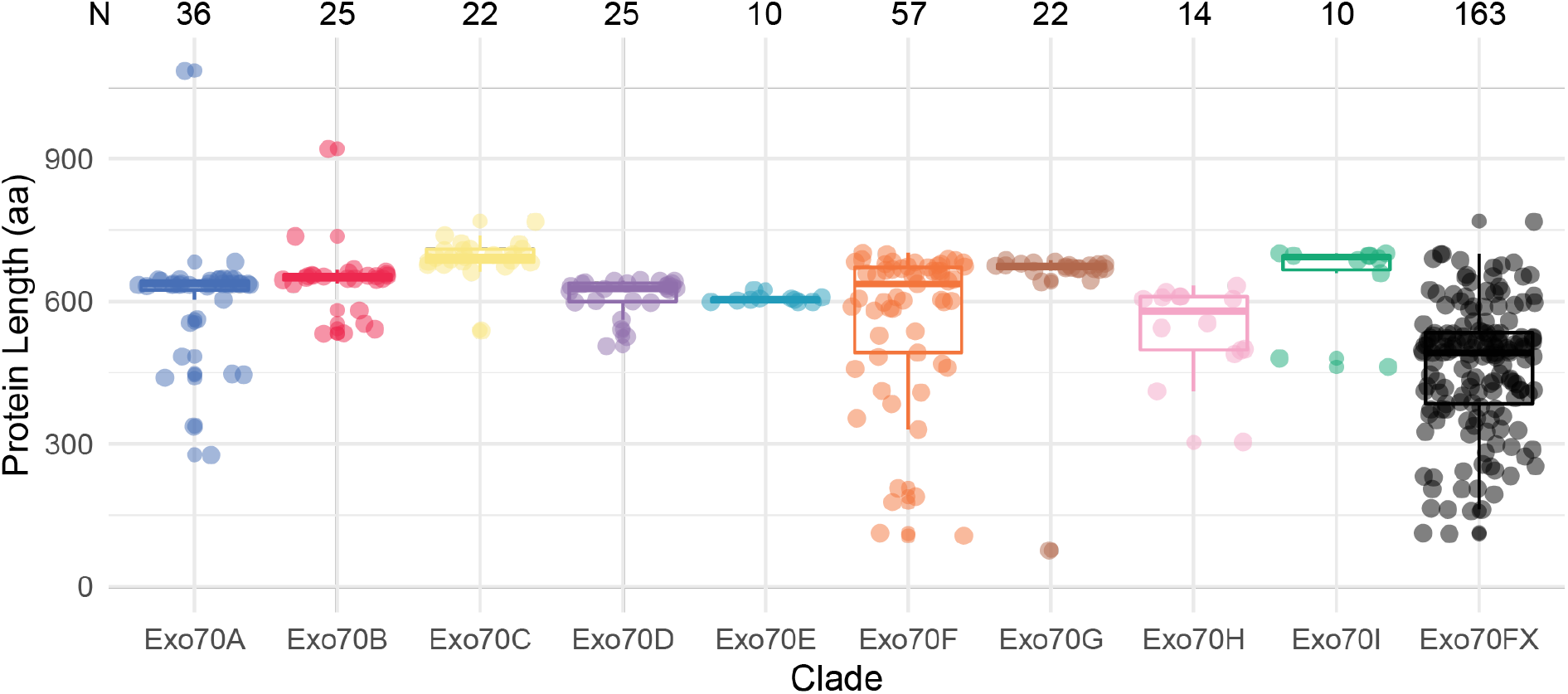
Members of the Exo70F and Exo70FX clades are experiencing a reduction in protein length. The y-axis is the length of predicted protein sequences (number of amino acids). Exo70 proteins from barley (*H. vulgare*), wheat (*T. aestivum*), purple false brome (*B. distachyon*), rice (*O. sativa*), *O. thomaeum*, maize (*Z. mays*), foxtail millet (*S. italica*), and sorghum (*S. bicolor*). Boxplots showing the 25^th^, 50^th^, and 75^th^ quantiles of protein length for Exo70 clade members. Upper and lower whiskers extend to the largest and smallest values no further than 1.5 times the inter-quartile range. Individual lengths are shown using jitter. N denotes the number of proteins in each Exo70 clade.

**Supplemental Figure 11.**
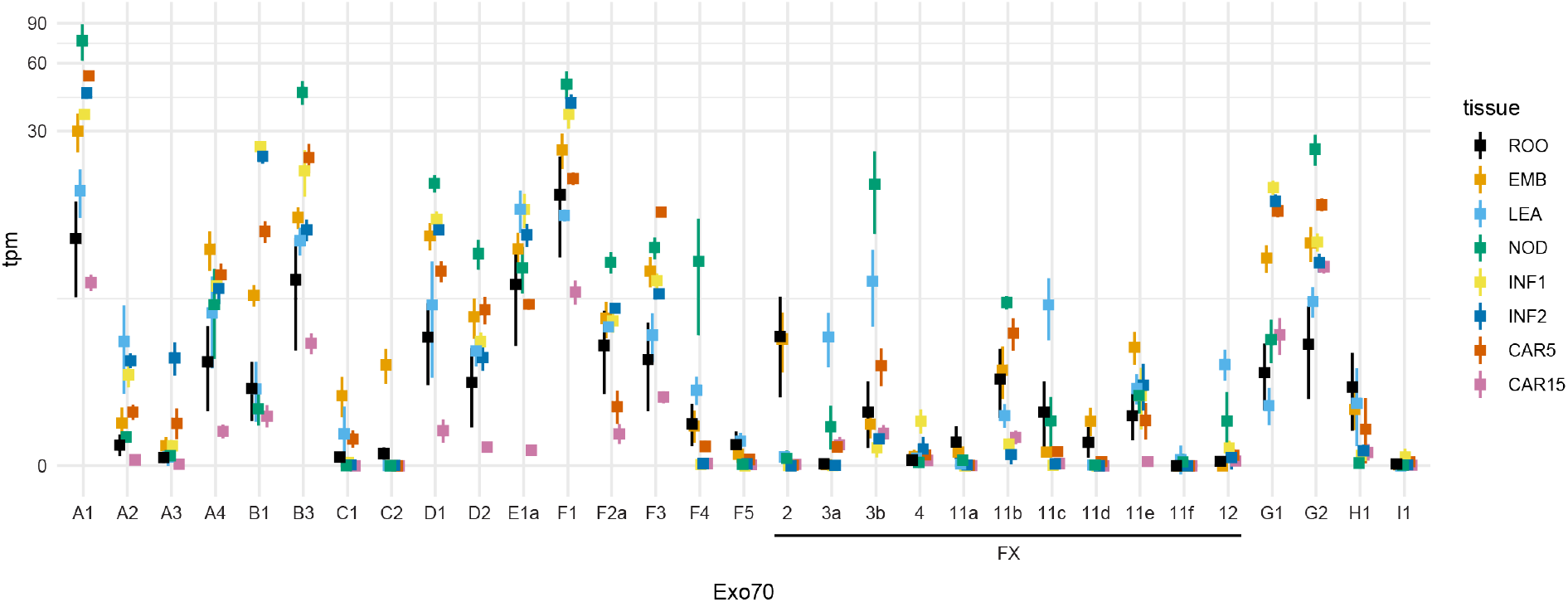
Expression analysis of barley Exo70 in eight diverse tissues. RNAseq data was obtained from EMBL/ENA accession ERP001600 (IBGSC, 2012), trimmed, and used to estimate expression level (transcripts per million (tpm); y-axis; pseudo-log scale) based on the barley predicted transcriptome (Mascher et al., 2021). Identifiers (x-axis) correspond to the suffix of Exo70 encoding genes. Mean and standard deviation are shown as a square and whiskers. Color coding and order shows the tissue including early developing grain (15 days post anthesis (CAR15) and 5 (CAR5)), germinating grain (4 day) embryos (EMB), early developing inflorescences (5 (INF1) & 15 mm (INF2)), shoots from seedlings (LEA; 10 cm stage), developing tiller internodes (NOD) (six-leaf stage), and roots from seedlings (ROO; 10 cm stage).

**Supplemental Figure 12.**
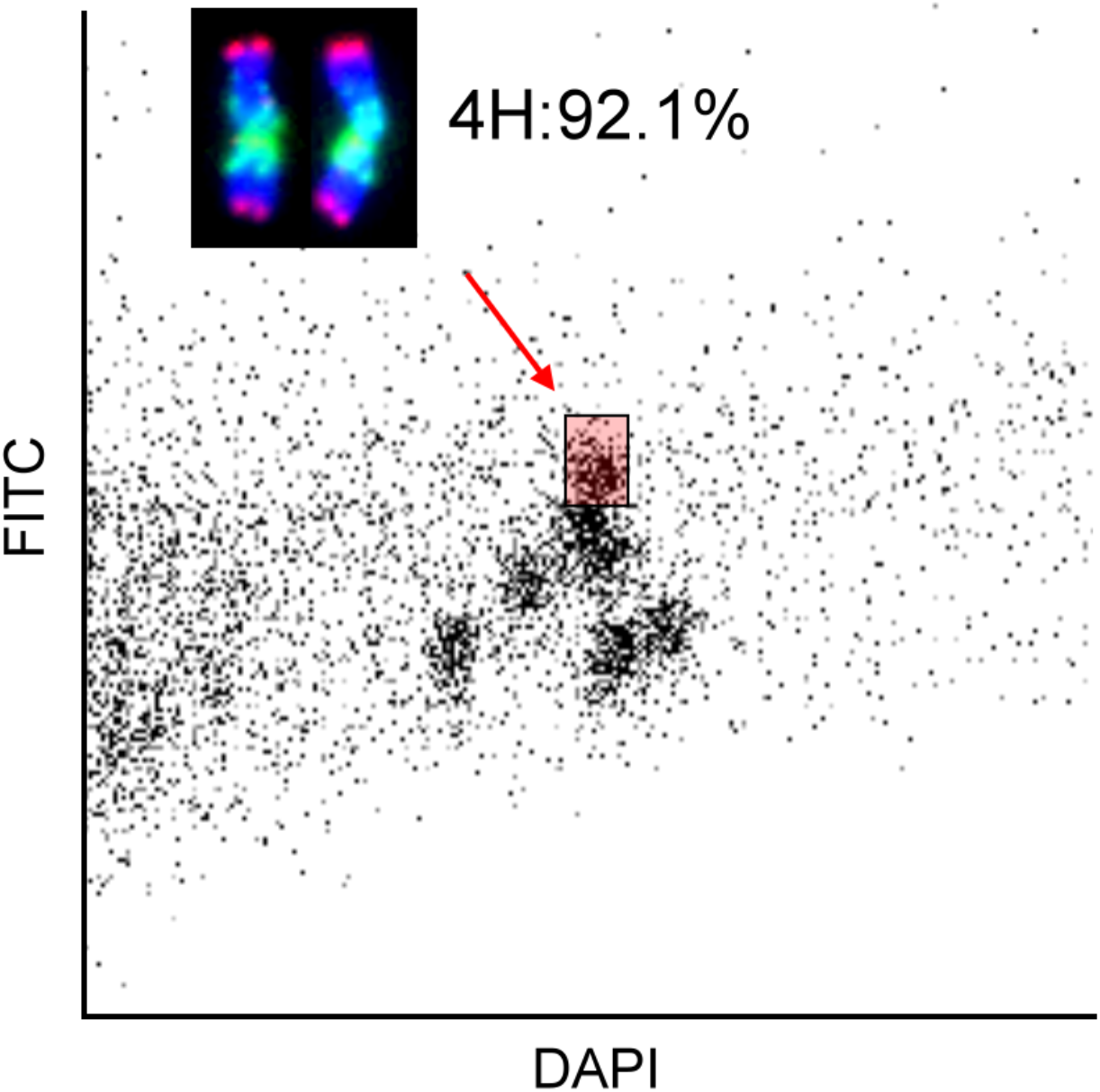
Bivariate flow karyotype DAPI vs. GAA-FITC obtained after flow cytometric analysis of mitotic metaphase chromosomes isolated from barley accession CI 16139. Barley 4H chromosomes were flow sorted at purity of 92.1% from the sorting window shown as red rectangle. Inset: Images of chromosome 4H flow-sorted onto a microscope slide. The chromosome was identified based on fluorescent labelling after FISH with probes for GAA microsatellite (green) and HvT01 satellite (red). The chromosomes were counterstained by DAPI (blue).

## Notes

https://doi.org/10.6084/m9.figshare.17288801

### Competing Interest Statement

The authors have declared no competing interest.

